# Sequelae and reversal of age-dependent alterations in mitochondrial dynamics via autophagy enhancement in reprogrammed human neurons

**DOI:** 10.1101/2025.10.02.680077

**Authors:** Eva Klinman, Ji-Sun Kwon, Roland E. Dolle, Stephen C. Pak, Gary A. Silverman, David H. Perlmutter, Andrew S. Yoo

**Affiliations:** Department of Neurology, Washington University School of Medicine, St. Louis, MO 63110, USA; Department of Developmental Biology, Washington University School of Medicine, St. Louis, MO 63110, USA; Program in Computational and Systems Biology, Washington University School of Medicine, St. Louis, MO 63110, USA; Department of Biochemistry and Molecular Biophysics, Washington University School of Medicine, St. Louis, MO, USA; Department of Pediatrics, Washington University School of Medicine, St. Louis, MO, USA; Department of Developmental Biology, Washington University School of Medicine, St. Louis, MO 63110, USA; Center of Regenerative Medicine, Washington University School of Medicine, St. Louis, MO 63110, USA

## Abstract

How aging of human neurons affects dynamics of essential organelle such as mitochondria and autophagosomes remains largely unknown. MicroRNA-induced directly reprogrammed neurons (miNs) derived from adult fibroblasts retain age-associated signatures of the donor, enabling the study of age-dependent features in human neurons, including longitudinal isogenic samples. Transcriptomic analysis revealed that neurons derived from elderly individuals are characterized by gene expression changes associated with the regulation of autophagosomes, lysosomes, and mitochondria, compared to young counterparts. To clarify these changes at the cellular level, we performed live-cell imaging of cellular organelles in miNs from donors of different ages. Older donor miNs exhibit decreased mitochondrial membrane potential, which surprisingly co-occurs with a significant increase in mitochondrial fission and fusion events. We posit that the increased fission and fusion of mitochondria may reflect age-dependent compensation for impaired mitochondrial turnover, perhaps due to changes in autophagy. We subsequently identified a significant decrease in autophagosome acidification in neurons derived from individuals >65 years compared to younger donors, and a corresponding age-dependent reduction in neuritic lysosomes resulting in fewer lysosomes available to acidify autophagosomes. This age-dependent deficit in autolysosome flux was rescued by promoting autophagosome generation through TFEB, which also reversed the age-dependent increase in mitochondrial fission and fusion and improved mitochondrial health. Partial organelle recovery occurred after inducing mitophagy or inhibiting mitochondrial fission. Together, this work reveals a mechanism by which aging reduces autophagic flux secondary to a loss of neuritic lysosomes, resulting in in mitochondria-intrinsic mechanisms to avoid loss of energy production.

## Introduction

Aging is a universal process and understanding the molecular changes which occur with age is key to determining mechanisms to slow aging and identify aberrant age-dependent diseases. Neurons pose a particular problem for the study of aging due to their inaccessible nature and post-mitotic state. Animal models have contributed substantially to knowledge of aging processes in the body and the brain. However, animals do not capture the complex genetic variability nor the age range of humans, whose lifespans can exceed 100 years (Mertens *et al*., 2018; Pitrez *et al*., 2024). This difference in time scale is paramount for neurons which must last the life of the organism. To better model human-specific neuronal traits, neuronal differentiation of induced pluripotent stem cells (iPSCs) has provided a powerful platform for studying neurogenesis and neurodevelopmental disorders over the past decade. However, due to the dedifferentiation process involved in generating iPSCs, age-related epigenetic and cellular encoding is erased. As a result, neurons derived from iPSCs are functionally embryonic (Mertens *et al*., 2018; Pitrez *et al*., 2024; Horvath, 2013; Patterson *et al*., 2012). Thus, although iPSCs excel at capturing the heterogeneity of human genetics, they cannot serve as an effective model of brain aging without extensive manipulation (Wu *et al*., 2019). In the absence of primary neurons harvested from human donors of different ages, direct fate conversion of alternative cells into neurons offers an alternative approach for modeling neuronal aging.

Ectopic expression of neurogenic factors in fibroblasts allows for cell fate conversion directly into neurons without undergoing a pluripotent stage (Mertens *et al*., 2016). This conversion process retains both the genetic diversity and cellular age of the human donor (Huh *et al*., 2016; Mertens *et al*., 2015). In particular, brain-enriched microRNAs (miRNAs), miR-9/9* and miR-124, have been employed as neurogenic effectors to efficiently induce chromatin reconfiguration, which in combination with subtype-defining transcription factors generates specific disease-relevant neuronal subtypes (Cates *et al*., 2021; Lu *et al*., 2021; Victor *et al*., 2018; Yoo *et al*., 2011; Abernathy *et al*., 2017; Victor *et al*., 2014). The resulting miRNA-induced neurons (miNs) retain the age signature stored in pre-reprogrammed fibroblasts and can be used to model both aging of human neurons and late-onset neurodegenerative disorders including Huntington disease, primary tauopathies, and Alzheimer disease (Capano *et al*., 2022; Church *et al*., 2021; Huh *et al*., 2016; Sun *et al*., 2024) To this end, directly reprogrammed miNs enable the study of how aging contributes to the onset of neurodegeneration in human cells (Capano *et al*., 2022; Lee *et al*., 2024; Oh *et al*., 2022; Sun *et al*., 2024).

The miN system reflects a molecular snapshot of neuronal age, allowing for critical investigation of age-associated changes in the behavior of intracellular organelles which can only be evaluated in living neurons. Existing models in other organisms have demonstrated that distribution and function of autophagosomes, lysosomes, and mitochondria change with age (Aman *et al*., 2021; Grimm and Eckert, 2017; Lipinski *et al*., 2010; Tian *et al*., 2023), although this has not been studied in human neurons. Arrangement and behavior of organelles depends on organization by the neuronal cytoskeleton, whose associated proteins vary substantially between animal models and humans. Similarly, an age-dependent decrease in autophagic function and biogenesis has been observed in animal neurons and non-neuronal human cells (Ott *et al*., 2016; Kaushik *et al*., 2012; Stavoe and Holzbaur, 2020), but has yet to be assessed in human neurons, although proteomics from a transdifferentiated human neuron model of Alzheimer’s disease show a deficit in lysosomal homeostasis (Chou *et al*., 2025). Decreased macroautophagy/autophagy drives changes in mitochondrial health, as autophagosome flux plays a crucial role in eliminating damaged and dysfunctional mitochondria (Lipinski *et al*., 2010). Expression profiling from aged human brains has implicated mitochondrial proteins and reactive oxygen species generated by mitochondrial respiration in the pathogenesis of aging (Fang *et al*., 2019), and mitochondrial dysfunction is posited to drive pathologic changes in Alzheimer dementia (Birnbaum *et al*., 2018; Chakravorty, Jetto and Manjithaya, 2019; Hawkins and Duchen, 2019). Existing work in other direct human neuronal differentiation models confirm that expression of mitochondrial genes related to oxidative phosphorylation and energy production are reduced with aging (Kim *et al*., 2018) and highlight the power of direct neuronal conversion to enable the study of mitochondrial health and glycolysis during aging compared to embryonic iPSC models (Varghese *et al*., 2025).

In this study, we investigated the behavior of autophagosomes and other crucial organelles in human neurons to evaluate systemic age-associated changes. We identified increased expression of RNA transcripts responsible for autophagosome regulation and transport with advancing age, and a simultaneous reduction in lysosomal and mitochondrial protein expression with age. We observed a loss of autophagosome acidification in the neurites from old donors, which corresponded to a decrease in neuritic lysosomal concentration. The flux deficit in autophagosomes could be rescued by chemically increasing autophagosome formation. This age-dependent decline resulted in a novel compensatory increase in mitochondrial quality control through the dynamic processes of fission and fusion, rather than reliance on degradation through mitophagy. Analysis of longitudinally collected samples suggests that these age-associated cytostatic changes occur around or before age 65 but after age 50. Excitingly, we demonstrated that inducing autophagy by activating the TFEB pathway was sufficient to rescue age-related mitochondrial reliance on fission and fusion without compromising mitochondrial health, suggesting a druggable pathway to combat this manifestation of aging. Overall, this work delineated a pathway by which aging reduced autolysosome flux due to a decrease in neuritic lysosomes, resulting in mitochondria which more readily interacted with each other to combat the loss of functional mitophagy. These changes took effect prior to age 65 corresponding with the onset of many sporadic neurodegenerative conditions including both Alzheimer and Parkinson diseases.

## Results

### Visualization of neuronal reprogramming process

The human fibroblasts used for neuronal reprogramming were sourced from a variety of biobanks and cover the range of the human lifespan, from newborns to advanced age. Donor lines were divided by age (**Table 1**), including neonatal (newborn foreskin-derived dermal cell lines, **Figure S1A**), young (age 20-35, **Figure S1B**), and old (age >75, **Figure S1C**). We implemented the established microRNA-mediated direct neuronal reprogramming protocol based on virally transduced microRNAs miR-9/9*-124 to transition fibroblasts to a neuronal fate (Church *et al*., 2021; Yoo *et al*., 2011). We included expression of dominant negative p53 (p53DD) to enhance reprogramming speed in the postmitotic reprogramming state (Cates *et al*., 2025) and transcription factors neuronal differentiation 2 (NEUROD2) and myelin transcription factor 1-like (MYT1L) to promote cortical lineage differentiation (**Figure 1A**). Fibroblasts were directly differentiated into neurons, which was reflected by the increased expression of the synapsin promoter (Cates *et al*., 2025) and outgrowth of dendrites and axons by post-infection day (PID) 20 (**Figure 1B**). By PID 30 miNs are phenotypically and transcriptionally mature (Cates *et al*., 2025). This process includes a shift in morphology mirroring the changing transcriptional landscape, from fibroblasts characterized by large flat spindle-shaped cells to neurons characterized by long thin projections radiating out from a central cell body (**Video V1)**. During the process of differentiation into cortical neurons, protein expression patterns changed to reflect the new identity of the cells, including extensive neurite outgrowth marked by the morphological microtubule marker beta-3 tubulin (TUBB3) and neuritic localization of the neuronal-specific microtubule-associated protein tau (MAPT/tau) (**Figure 1C**). All selected fibroblast lines regardless of donor age reliably reprogrammed into neurons (92.5-96.4% of cells), as determined by expression of the neuron-specific protein tau and cellular morphology (**Figure S1D**), consistent with the previous reports (Cates *et al*., 2021; Cates *et al*., 2025; Sun *et al*., 2024).

**Table 1.** Identity and source for patient fibroblast samples. ATCC = American Type Culture Collection biobank Coriell = NIH-associated biobank DIAN = Dominantly Inherited Alzheimer Network research center biobank

| ID | SYMBOL | AGE | GENDER | DISEASE | SITE | SOURCE |
| --- | --- | --- | --- | --- | --- | --- |
| neoHDF | ● | neonatal | Male | Healthy | Foreskin | ATCC |
| HDF-F | ■ | neonatal | Male | Healthy | Foreskin | ScienCell |
| AG04054 | ● | 29 | Male | Healthy | Arm | Coriell |
| AG08790 | ■ | 31 | Male | Healthy | Arm | Coriell |
| AG05411 | ▲ | 33 | Male | Healthy | Arm | Coriell |
| GM02171 | ● | 22 | Female | Healthy | Unknown | Coriell |
| F12-442 | ■ | 30 | Female | Healthy | Hip | DIAN |
| AG09599 | ▲ | 30 | Female | Healthy | Arm | Coriell |
| AG13076 | ● | 82 | Male | Healthy | Arm | Coriell |
| AG14036 | ■ | 84 | Male | Healthy | Arm | Coriell |
| AG13129 | ▲ | 89 | Male | Healthy | Arm | Coriell |
| AG11246 | ● | 78 | Female | Healthy | Arm | Coriell |
| AG08269 | ■ | 82 | Female | Healthy | Arm | Coriell |
| AG13077 | ▲ | 85 | Female | Healthy | Arm | Coriell |

**Figure 1:**
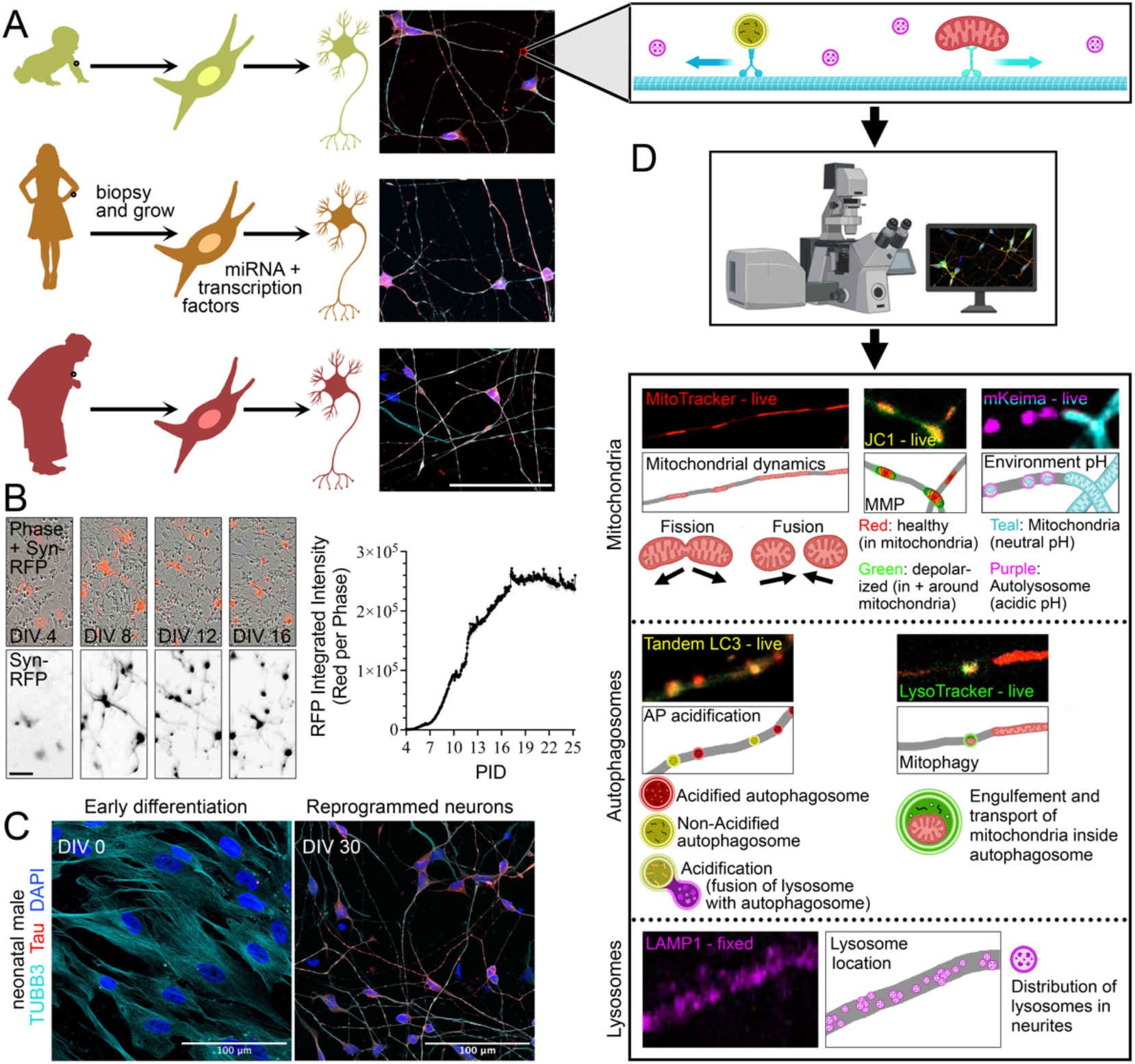
Neuronal conversion of young and old fibroblasts enables live-cell organelle imaging. **(A)** Illustration of miN direct differentiation process, wherein skin biopsies from individuals of different ages are cultured for fibroblasts and differentiated using miRNA and transcription factors into miNs that maintain characteristics of the age of the donor; donor fibroblasts generate miNs with high efficiency at all donor ages (right). **(B)** Still frame images from time-lapse video of differentiating cells expressing pSynapsin-RFP (left), quantification of RFP intensity over region containing reprogrammed neurons in phase contrast over the course of neuronal reprogramming (right). Full video available as Supplemental Video 1. **(C)** Immunofluorescence representative images of the transition from fibroblasts (DIV 0) to mature (DIV 28) in a neonatal donor. Immunostaining was performed for neuronal markers TUBB3 and tau, as well as DAPI. **(D)** On top, schematic of evaluation for organelle characteristics in living miNs including motor proteins (teal) transporting an autophagosome (yellow) and mitochondria (red) in the presence of lysosomes (purple). On bottom, overview of imaging approaches including mitochondrial, autophagosome, and lysosomal assessment. Scale bar 100 µm in A, B, and C.

### Expression of organelle-associated transcripts change with age

Dysregulation of neuronal organelles, particularly autophagosomes and mitochondria, have been linked to the pathogenes -is of aging and are tied to the pathologic changes associated with neurodegenerative conditions such as Alzheimer disease (Fang *et al*., 2019; Chakravorty, Jetto and Manjithaya, 2019; Birnbaum *et al*., 2018; Swerdlow, 2018; Nixon *et al*., 2005; Lipinski *et al*., 2010). Many of the features of organelle behavior and function rely on the use of living cells, which had made them difficult to assess in human neurons. As such, we sought to establish a model of intracellular organelle behavior during neuronal aging. We began by performing analysis of an RNA sequencing data set previously collected from cortically differentiated miNs from young and old healthy donors (Sun *et al*., 2024). Pathway analysis of differentially expressed gene (DEG) lists using PANTHER indicated notable biological and cellular processes which increased or decreased with age (Chen *et al*., 2013; Kuleshov *et al*., 2016; Xie *et al*., 2021). Moreover, advancing age resulted in reduced expression of transcripts associated with mitochondrial function (**Figure S2A**), consistent with prior studies from human post-mortem brains including decreased mitochondrial transcripts with age (Herring *et al*., 2022; Jeffries *et al*., 2025). In contrast, DEGs associated with regulation of autophagy, mitochondrial distribution and kinetics, and transport of organelles increased with aging (**Figure S2B**). Notably, mitochondrial transcripts were among the most significantly decreased gene ontology terms as a function of age, while many of the top upregulated transcripts involved neuron maturation (**Figure S2C-D**).

To extend these findings and study the behavior of organelles identified by transcriptome analysis in living neurons from donors of different ages, we used live-cell imaging and functional assessment to investigate the age-related change in mitochondria, autophagosome, and lysosomes in miNs derived from individuals across the human lifespan. Transcripts involved in different mitochondrial pathways were markedly decreased with age, thus we began our assessment of organelles in living neurons by evaluating mitochondria. We evaluated the behavior and function of live mitochondria using the organelle marker MitoTracker, membrane readout JC-1, and mitophagy marker mt-mKeima, quantified the acidification and transport of autophagosomes using pH sensitive tandem-LC3 and organelle maker LysoTracker, and assessed lysosomal distribution using immunofluorescence (**Figure 1D**).

### Live mitochondria exhibit signs of age-related stress in neurons

Mitochondria are particularly important to neurons which require high production of ATP to maintain their membrane potential, release neurotransmitters, and propagate action potentials (Rangaraju *et al*., 2019). Mitochondrial health is primarily maintained by two biological processes: mitophagy and mitochondrial fission or fusion. Mitophagy is the well-characterized selective breakdown of damaged mitochondria by autophagosomes (Stavoe and Holzbaur, 2020; Toescu, Myronova and Verkhratsky, 2000). The second mechanism of mitochondrial preservation involves merging with other mitochondria to facilitate the exchange of membrane, proteins, and DNA in a process called mitochondrial fusion, or dividing into new smaller mitochondria in a process termed fission (Liu *et al*., 2020; Zhang *et al*., 2019). Thus, ailing mitochondria can be partially rescued by exchange of components with their neighbors. Between these processes, neurons can maintain mitochondrial health and function to avoid loss of ATP production or the dangerous generation of free oxygen radicals.

We found that mitochondria in miNs were spread throughout the neurites and cell body as expected in neurons (Pekkurnaz and Wang, 2022), which contrasted with the contiguous networked web of mitochondria observed in fibroblasts (**Figure 2A**). For live-cell reporting of the health of endogenous neuronal mitochondria, miNs were incubated with the dye JC-1, which provides a color-shifting fluorescent readout of mitochondrial membrane potential (MMP). When MMP is intact, JC-1 forms aggregates inside of mitochondria which fluoresce red, while when JC-1 dissociates into monomers in response to loss of MMP it fluoresces green in both mitochondria and in the cytoplasm (**Figure 2B**). Comparing the red:green ratio of JC-1 signal, we found that miNs derived from older donors showed more MMP disruption than miNs from younger donors (**Figure 2C**), consistent with prior work (Kim *et al*., 2018).

**Figure 2:**
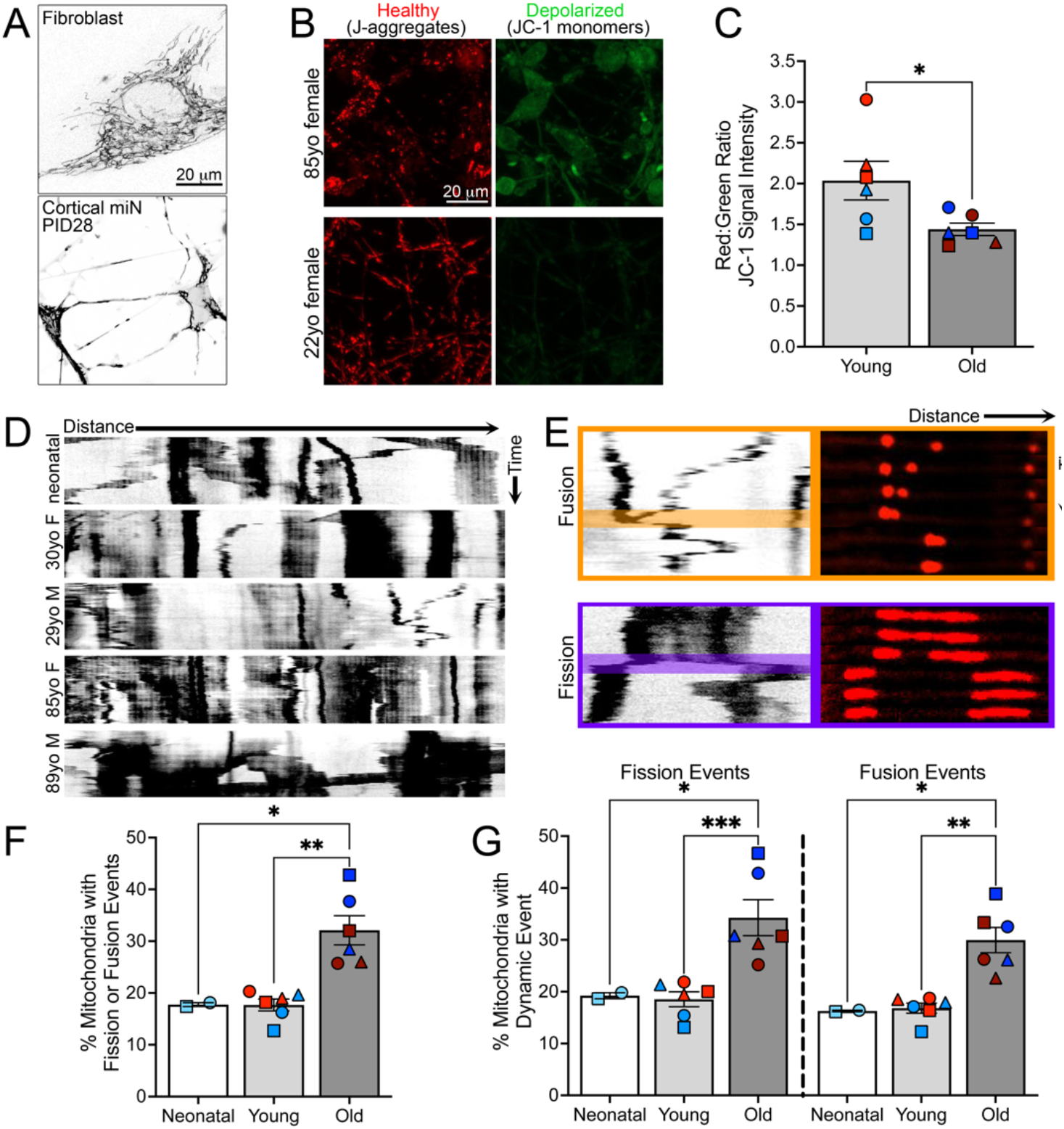
Neuronal mitochondrial characteristics change with advancing age. **(A)** Representative image of MitoTracker labeled mitochondria distribution during differentiation from fibroblasts (top) to cortical miNs (bottom). Scale bar 10µm. **(B)** Z-projection of live mitochondria labeled with JC-1 to identify intact mitochondrial membrane potential (red) and loss of membrane potential (green) in young (bottom) vs old (top) donor miNs. Scale bars 20 um. **(C)** Quantification of mitochondrial membrane potential in miNs from young and old donors as determined by JC-1 fluorescent assay. **(D)** Representative kymographs of Mito-Tracker-labeled mitochondria (black-on-white) from donors of different ages and sexes. **(E)** Examples of fusion (top, orange) and fission (bottom, purple) events in mitochondria, series of 6 still frames from the highlighted portion of the fusion and fission kymographs to the right to assist visualization. **(F)** Quantification of percent total mitochondria undergoing dynamic fission or fusion events in miNs from donors of different ages. **(G)** Fission and fusion events from F separated by event type and age of donor calculated by percent total mitochondria present. Each graph symbol represents a distinct donor, symbols correspond to donor identities in Table 1. Each donor underwent ≥2 reprogramming events totaling ≥3 independent wells and ≥6 neurites analyzed per donor. Mean ± SEM; t-test used for calculation in C, 2-way ANOVA in F-G; *p<0.05, **p<0.01, ***p<0.001

To determine what age range these changes occur in more detail, we turned to isogenic fibroblast samples collected years apart from the same individuals. The ability to use longitudinal fibroblast samples has previously provided insights into age-related neuronal changes in patients with Huntington disease (Lee *et al*., 2024). We identified 6 donors’ fibroblast samples collected approximately 15 years apart available through the Baltimore Longitudinal Study of Aging. Of note, all identified donors were male as no corresponding female longitudinal samples were available in the same age range. The identified donors provided a skin biopsy at approximately age 30 and again in their late 40s, or in their late 60s and again in their mid-80s (**Figure S3A**). miNs were generated from these longitudinally collected samples (**Figure S3B**), and we found that mitochondrial membrane potential did not differ over 15 years within isogenic lines, while there appeared to be an inflection point prior to age 65 between the young and old donor sets. Prior to age 50, neurons derived from fibroblasts behaved ‘young,’ while beyond age 65, the observed changes in mitochondrial health were present and static (**Figure S3C**). As such, we were comfortable continuing with our evaluation of the youngest and oldest donors in our fibroblast cohort.

Our analysis of existing RNA sequencing results additionally identified an age-dependent increase in expression of genes associated with transport of mitochondria (**Figure S2B**), which is required for mitochondria to encounter neighboring mitochondria. To determine if interactions between neighboring mitochondria in neurites alters with age, we assessed the dynamic fission and fusion of native unperturbed mitochondria in live miNs using MitoTracker Red CMXRos (MitoTracker) over time by kymographs (**Figure 2D, Video V2**). Fission and fusion were apparent when mitochondria either separated from a single mass or joined together during the recording (**Figure 2E** and **Video V3)**. We quantified the percent of mitochondria in the field of view which underwent dynamic fission or fusion events during the 5-minute live-cell recording and confirmed that both fission and fusion increased significantly with advancing ages in miNs (**Figure 2F**). Surprisingly, there was no clear preference for fission or fusion and both events occurred in an even ratio (**Figure 2G**). Although changes in mitochondrial fission and fusion have been observed in aging in non-human models, results vary widely between studies (Liu *et al*., 2020). We confirmed this shift in fission and fusion events in our longitudinal samples and again found a cutoff prior to age 65 (**Figure S3D**) dividing our young and old donor sets. Intriguingly, this shift in mitochondrial behavior corresponds to the age at which individuals experience an ever-increasing risk of developing sporadic neurodegenerative disorders such as Alzheimer disease (Liang *et al*., 2021). Our findings imply that aged human neuronal mitochondria may act to compensate for mitochondrial defects such as decreased MMP through increased mixing of mitochondrial components.

Other mitochondrial characteristics such as percent motility, length, or density along the neurite were unaffected by aging. Motile fraction remained stable at 30-40% (**Figure S3F**), consistent with prior work in animal models and other human cells (Misgeld and Schwarz, 2017), and length (**Figure S3G**) likewise fell in the range established for neuritic mitochondria in fixed human brain tissue and living rat neurons (Kim *et al*., 2022; Rangaraju, Lauterbach and Schuman, 2019). Interestingly, analysis of mitochondria in the brains of living mice indicated more dramatic reduction of both motility and length of mitochondria with age (Lewis *et al*., 2016). Of note, neurites with motile mitochondria may be enriched in this work because individual neurites are easier to identify when mitochondria traverse their length; as such, non-selected neurites may have a lower motile fraction. Mitochondrial density also remained unchanged across conditions (**Figure S3H**), at approximately 15 mitochondria per 100 µm of neurite. Of note, this value differs from measurements taken in other animals, which vary between 5 (*C. elegans*) and 50 (rat) mitochondria per 100 µm (Mondal *et al*., 2021; Sure *et al*., 2018; Wareski *et al*., 2009), highlighting the importance of studying human neuronal processes in addition to animal models.

A crucial pathway for controlling the degradation of dysfunctional mitochondria is mitophagy, the selected autophagy of mitochondria. Our analysis of existing RNA sequencing data indicated a significant increase in expression of transcripts associated with autophagosomes, as well as a decrease of those associated with mitochondrial function (**Figure S2A-B**). Given the results of our mitochondria data, we sought to determine if increased mitochondrial fission and fusion events with aging might represent changes to compensate for a decline of autophagy function in neurons.

### Neurons from old donors have functional autophagosome deficit

Autophagosomes form at the terminus of neurites (the axon and dendrites) or along their length, and transport towards the cell body collecting debris for breakdown (Maday, Wallace and Holzbaur, 2012). The functional lysis of internalized debris depends on acidification of the autophagosome to form an autolysosome (Mauvezin *et al*., 2015). To evaluate functional acidification of autophagosomes into autolysosomes, we implemented viral expression of a tandem labelled construct (Pankiv *et al*., 2007). Microtubule-associated protein 1A/1B-light chain 3 (LC3), is expressed on the inner membrane of autophagosomes and is exposed to the internal milieu of the organelle including its pH. The tandem LC3 construct contains both mCherry and GFP fluorophores (**Figure 3A**). The mCherry construct is acid-stable to a pH of 4.0 and continues to fluoresce in the pH 4.8 environment present during autolysosome acidification (Pankiv *et al*., 2007). In contrast, the GFP tag is quenched at a pH of approximately 6.0. Thus, the tandem LC3 construct is capable of distinguishing non-acidified autophagosomes from their acidified autolysosome siblings using a color-shift readout, in addition to providing information on autophagosome quantity.

**Figure 3:**
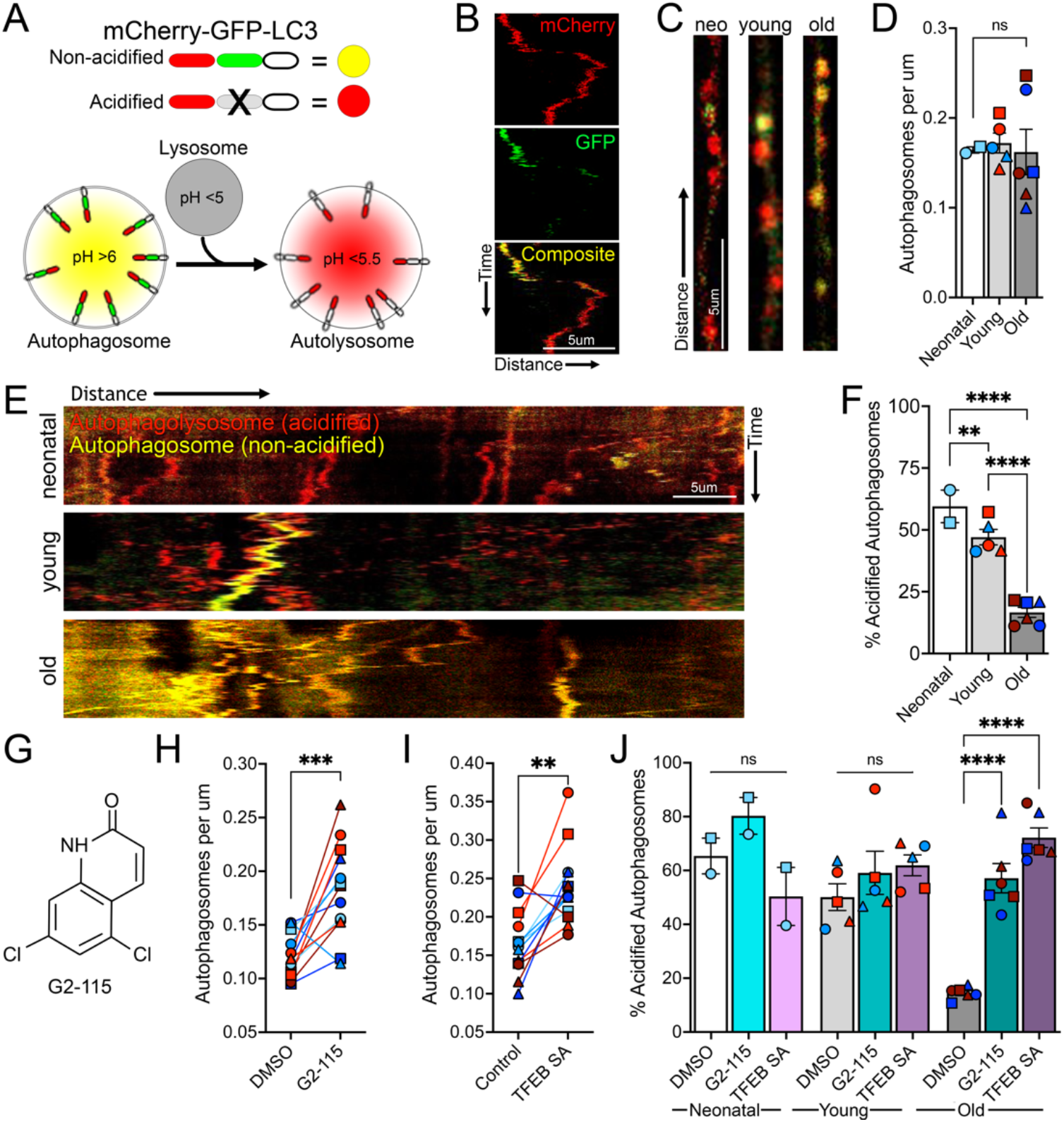
Autolysosome formation decreases with donor age but recovers in response to transcription-level stimulation in cortical miNs. **(A)** Tandem LC3 construct includes acid-stable mCherry tag, acid-quenched GFP tag, and autophagosome marker LC3. When exposed to an acidic environment via fusion with a lysosome, GFP fluorophore is lost, resulting in construct color change from yellow to red. **(B)** Kymograph of an autophagosome becoming acidified with quenching of GFP signal halfway through recording. **(C)** Straightened segments of neurites demonstrating acidified (red) or non-acidified (yellow) autophagosomes in miNs from donors of different ages. **(D)** Density of autophagosomes per micron in neurites from donor miNs of different ages and sexes. **(E)** Representative kymographs of autophagosomes in cortical miNs of different donor ages expressing tandem LC3 construct. Kymographs arranged such that presumed cell body is to the left, consistent with retrograde movement of autophagosomes in neurites. **(F)** Quantification of percent total acidified autophagosomes. **(G)** Chemical structure for autophagy inducer G2-115. **(H)** Quantification of density of autophagosomes (acidified and non-acidified) in miNs from the same donor treated with DMSO or G2-115. **(I)** Quantification of density of all autophagosomes expressing obligate nuclear TFEB SA. **(J)** Quantification of acidified autophagosomes in miNs treated with DMSO, G2-115, or TFEB SA from the same donors across different ages. Scale bar 5 µm in B, C, and E. Each donor line differentiated at least twice, ≥6 neurites collected from ≥3 independent wells per donor line. Mean ± SEM; 2-way ANOVA; **p<0.01, ***p<0.001, ****p<0.0001

Expression of the tandem LC3 construct in miNs does not impact cell vitality, and miNs continue to stably express the construct for at least 6 days (Lee *et al*., 2024). We used this assay in living neurons to assess in real-time the behavior, distribution, and acidification of autophagosomes (**Video V4**). Recordings were converted into two-dimensional kymographs to permit easy analysis and comparison of autophagosome density, motility, and acidification (**Figure S4**). Individual autophagosomes which acidified during the recording changed from yellow to red, although this rare event was observed less than 10 times in over 26 hours of recordings (**Figure 3B**). Traces from individual neurites were used to assess autophagosome density and distribution (**Figure 3C**). Autophagosome density remained stable at approximately 0.18 organelles per micron in all 14 donor lines (Fig. 3D), consistent with previous work in cultured mouse neurons, although murine work revealed decreased autophagosome biogenesis with age (Tsong, Holzbaur and Stavoe, 2023). As expected, movement of autophagosomes was predominantly unidirectional, consistent with known transport of autophagosomes from the distal axon towards the cell body (**Figure 3E**) (Klinman and Holzbaur, 2016). The location of an individual neurite’s cell body could not be conclusively determined due to the density of miNs required to maintain cell health (**Figure S1**) but neurites with clear associated soma or terminals within the imaging frame were not selected for analysis to enrich for analysis of the mid-neurite region.

Autophagosomes in miNs derived from newborn and young donors exhibited a mix of acidified and non-acidified autophagosomes (59.50% acidification in neonates and 47.06% in young). In contrast, miNs derived from old donors had an overwhelming majority of non-acidified autophagosomes reflecting a significant change with age (16.64% acidification) (**Figure 3F**). We observed a shift from mostly acidified vesicles in the young neuron group to predominantly non-acidified in the old neuron group, but not within the isogenic paired samples (**Figure S3E**). Our findings demonstrated that acidification of autophagosomes in neurites decreased with aging in human neurons, while the density of autophagosomes remained constant.

### Chemical induction of autophagy rescues age-dependent loss of autolysosome biogenesis

To determine if decreased autolysosome formation with age could be rescued by increasing autophagy, we employed a small molecule, G2-115 (**Figure 3G**), which was previously shown to increase autophagy in human neurons (Lee *et al*., 2024; Oh *et al*., 2022). G2-115 is a derivative of the sulfonylurea drug glibenclamide and contains pro-autophagic function (Hidvegi *et al*., 2010; Wang *et al*., 2019).

We cultured miNs derived from donors of different ages with G2-115 or solvent control (dimethyl sulfoxide, DMSO) to determine if enhancing autophagy during neuronal maturation would improve formation of autolysosomes. Neither G2-115 nor DMSO interfered with neuronal maturation (Lee *et al*., 2024). G2-115 significantly increased the overall density of autophagosomes in neurites by 60% across all age groups (**Figure 3H**). However, acidification of autophagosomes remained stable in the G2-115-treated miNs in the neonatal and young cohort, but improved significantly in the old donor miNs in response to G2-115 (**Figure 3J**, grey and teal bars). These results confirm that the age-dependent deficit in autolysosome flux could be rescued by promoting autophagosome formation using G2-115.

To better assess how G2-115 improves autophagosome acidification, we evaluated its mechanism of action. The G2 compound was found to decrease the interaction between RCAN1 and calcineurin, a calcium-and-calmodulin-dependent serine/threonine phosphatase (Lee *et al*., 2024). When freed from inhibition by RCAN1, calcineurin dephosphorylates its targets including the transcription factor TFEB (Klee, Crouch and Krinks, 1979; Rusnak and Mertz, 2000; Li *et al*., 2020). Dephosphorylated TFEB then enters the nucleus as a master regulator transcription factor to promote autophagy (Sardiello *et al*., 2009; Settembre *et al*., 2011; Settembre *et al*., 2013; Martina *et al*., 2012). To determine if nuclear TFEB is sufficient to recapitulate the effects of G2-115 on autophagosome behavior, we expressed obligately nuclear TFEB, in which the two calcineurin-mediated phosphorylation sites have been mutated from serine to alanine (TFEB S142/211A or TFEB SA) (**Figure S5A**) (Lee *et al*., 2024). Expression of TFEB SA through lentiviral transduction on PID 14 (the time point that aligns with G2-115 treatment) resulted in an increase in autophagosome density similar to that observed in miNs treated with G2-115 (**Figure 3I**). Likewise, TFEB SA reproduced the increased acidification of autophagosomes in miNs derived from aged donors without significantly changing the autophagosome population in younger donors (**Figure 3J**, purple bars). Together, this confirms that TFEB SA phenocopies G2-115, and that G2-115 improves autophagosome density and acidification via its effect on calcineurin and downstream nuclear localization of TFEB

We next sought to identify the upstream cause of the decreased autolysosome formation. We turned to lysosomes, as fusion of lysosomes with autophagosomes is required for the functional acidification of nascent autophagosomes into autolysosomes, and blocking this interaction results in cytotoxicity (Button *et al*., 2017).

### Aging is associated with lysosomal exclusion from neurites

Lysosomes, which are distributed throughout neurites, perform a variety of functions, including cell signaling and acidification of autophagosomes (Luzio, Pryor and Bright, 2007). Lysosomes provide the degradative power to autophagosomes via fusion, release of hydrolases, and an acidic environment (Ferguson, 2019; Luzio, Pryor and Bright, 2007). To date, the number of fusion events required to acidify an autophagosome as it is transported from the distal neurite to the cell body is unknown (Cason *et al*., 2022; Farfel-Becker *et al*., 2019), as is the distribution of lysosomes in human neurons as a function of age (Malik *et al*., 2019). Given their role in the acidification of autophagosomes, we evaluated if changes in lysosomal distribution may contribute to our findings of decreased autolysosome formation in neurites from older donor miNs.

We addressed the positioning of lysosomes in miNs derived from neonatal, young, and old donors through immunocytochemistry and confocal microscopy. Staining of mature cortical miNs of from donors of different ages demonstrated a dramatic change in the location of lysosomal associated membrane protein 1 (LAMP1)-positive lysosomes (**Figure 4A**). In neonatal and young donor miNs, lysosomes were readily apparent along neurites stained for tubulin, while miNs from old donors had lysosomes within the cell body but excluded from neurites (**Figure 4B**). This change in localization out of neurites with ageing was confirmed by both intensity of LAMP1 staining in neurites and Mander’s M1 correlation (**Figure 4C**). Previous work in other models of aging and age-associated diseases reported changes in lysosome acidification, but alterations in lysosomal distribution has never been described (Nixon, 2020).

**Figure 4:**
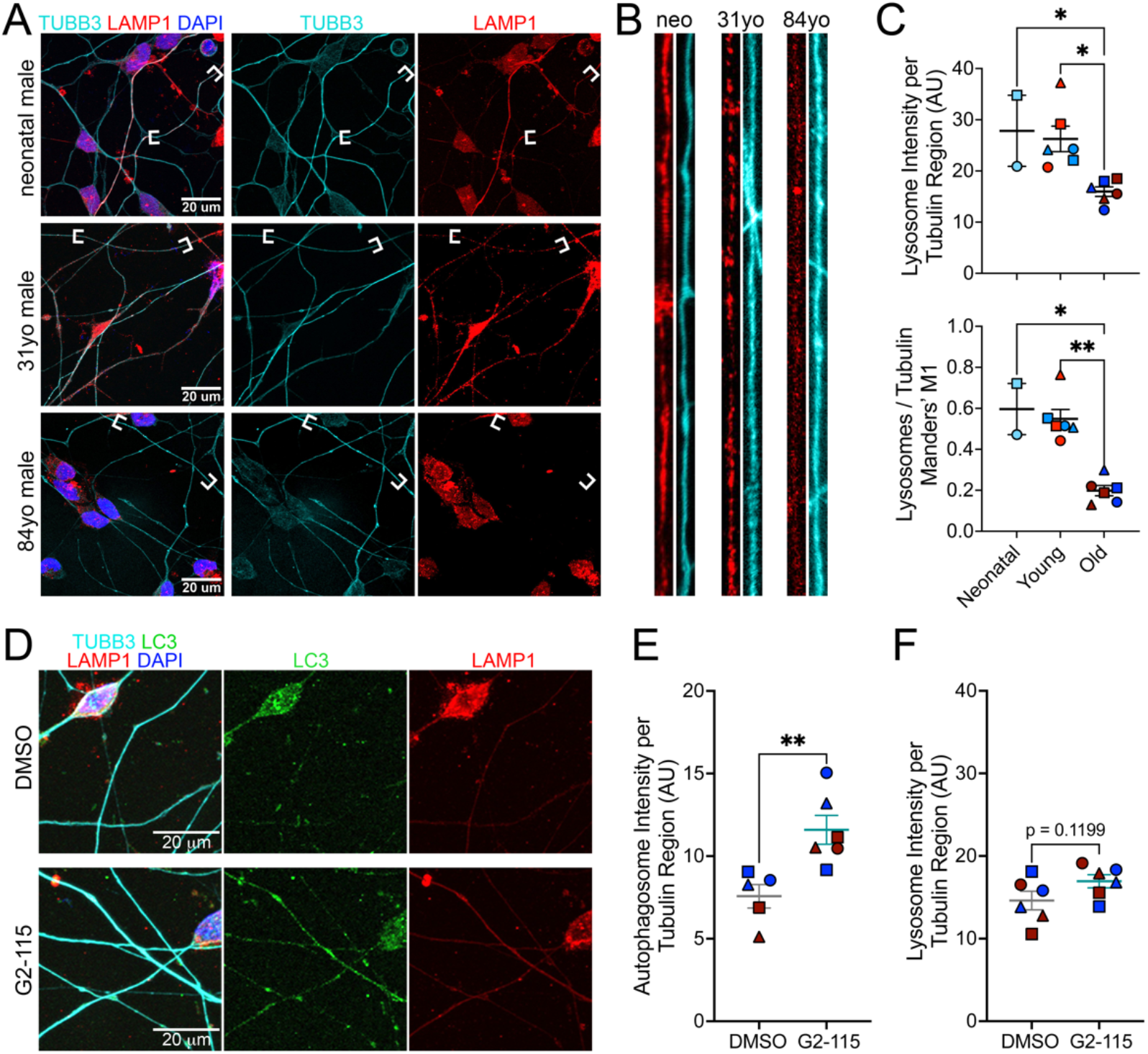
Neuritic-compartment lysosomes are lost with age and do not redistribute in response to increased autophagy in cortical miNs. **(A)** Representative immunofluorescence images of tubulin (TUBB3), lysosomes (LAMP1), and nuclei (DAPI) in stained cells from donors of different ages. White brackets highlight individual neurites magnified in B. **(B)** Straightened segments of neurites depicted between white brackets from A. Magnified neurites are immunostained for lysosomes (red, LAMP1) and microtubules (teal, TUBB3). Lysosome signal is decreased in miNs from older donors. **(C)** Lysosome signal intensity in tubulin-positive neurites (top) and fraction of tubulin which also contains lysosomal signal calculated using Manders’ M1 overlap coefficient (bottom) in neurites from different donors. **(D)** Representative immunofluorescence images of tubulin (TUBB3), autophagosomes (LC3), lysosomes (LAMP1), nuclei (DAPI) in stained cells from an 82-year-old female donor treated with DMSO or G2-115. **(E)** Autophagosome signal intensity in tubulin-positive neurites in miNs from donors treated with DMSO or G2-115. **(F)** Lysosome signal intensity in individual neurites calculated based on tubulin-positive regions as a function of treatment with DMSO or G2-115 in miNs derived from old donors. Scale bars 20µm. Two coverslips per done, 6 neurites per donor line collected over 3 fields of view per coverslip. Mean ± SEM; 2-way ANOVA used for quantification in C, unpaired t-test used for quantification in E and F; *p<0.05, **p<0.01

If the primary driver of decreased autolysosome flux is the inability to interact with lysosomes in neurites due to changes in lysosomal distribution, then the autolysosome rescue by G2-115 should alter lysosomal localization. To test this, we treated miNs from donors of all ages with G2-115 or DMSO and stained for LAMP1. Treatment with G2-115 did not significantly change the pattern of neuritic lysosomes in miNs from old donors, and lysosomes remained scarce within neurites (**Figure 4D**). Co-staining with LC3 confirmed an increase in autophagosome density in response to G2-115 treatment (**Figure 4E**). Quantification of lysosomal distribution confirmed that lysosome levels in the neurites remained surprisingly low despite treatment with G2-115, although there was a trend towards increasing lysosomes in neurites (**Figure 4F**). The reduction of neuritic lysosomes despite autolysosome improvement emphasizes the importance of redundancy in intracellular pathways responsible for autolysosome formation. These findings further support the theory that autophagosome acidification requires a low number of lysosomal fusions events (Cason *et al*., 2022), and suggests enhanced autolysosome formation as driving acidification in response to G2-115 rather than improved lysosomal distribution.

### Chemical rescue of autophagy improves mitochondrial health and function in aged neurons

Our results so far indicate that advancing age led to accumulation of damage to mitochondria, resulting in decreased mitochondrial membrane potential in neurons, and a compensatory increase in fission and fusion to distribute damaged mitochondrial components. We further determined that aging causes a defect in autophagosome acidification, which could be rescued via induction of autophagy using G2-115 or direct activation of the calcineurin-TFEB pathway. To address whether the autophagy defect may cause the observed change in living neuronal mitochondria, we evaluated if increasing autophagy using G2-115 or TFEB SA would restore markers of mitochondrial health including membrane potential and dynamic fission and fusion events to levels observed in healthy young individuals.

Using miNs generated from our 6 donors over the age of 65, we added DMSO or G2-115 to the culture media for 2 weeks or virally introduced TFEB SA and again imaged changes in MMP using JC-1. Surprisingly, we found that the addition of G2-115 or TFEB SA was sufficient to improve MMP to the level observed in miNs generated from young donors (**Figure 5A**), thus resetting the mitochondrial health through intervention on the autophagy pathway.

**Figure 5:**
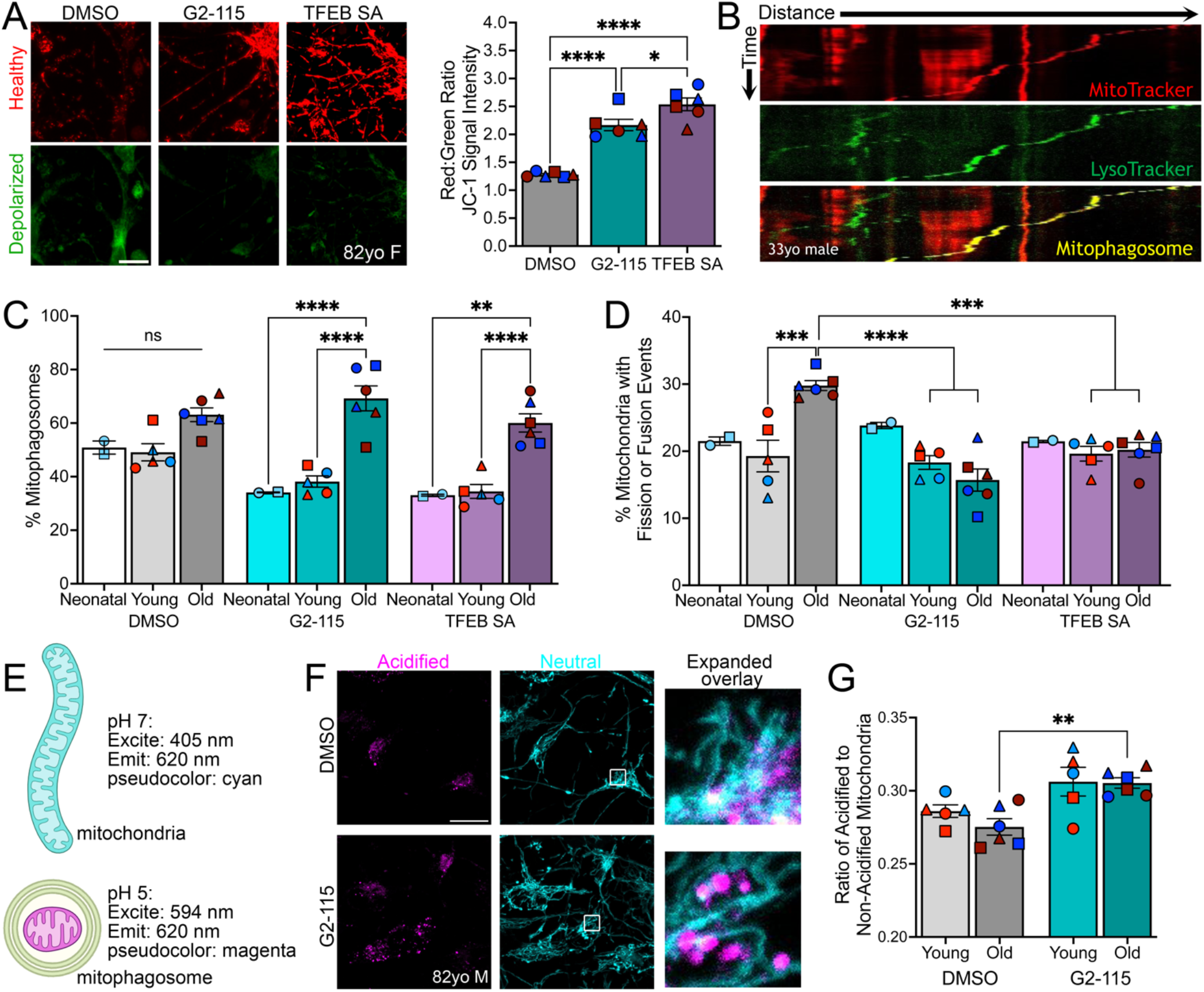
Age-dependent changes in mitochondrial behavior and health in response to autophagy enhancement. **(A)** Z-projection of live mitochondria labeled with JC-1 to identify healthy intact mitochondria (red) and those with loss of membrane potential (green) in DMSO-treated (left), G2-115-treated (middle), and TFEB SA-treated (right) miNs. Quantification from 6 old donors on the right. **(B)** Kymograph of MitoTracker (red) labeled mitochondria and LysoTracker (green) labeled autolysosomes in miNs. Mitophagosome (yellow) visible in overlay. **(C)** Quantification of percent of autophagosomes containing mitochondrial fragments by age and sex of donor in miNs treated with DMSO, G2-115, or TFEB SA. **(D)** Quantification of percent fission and fusion events of mitochondria in miNs from donors of different ages treated with DMSO, G2-115, or TFEB SA. **(E)** Schematic of mt-mKeima demonstrating different excitation wavelengths for neutral (pH 7, top, teal) and acidified (pH 5, bottom, magenta) mitochondrial compartments. **(F)** Z-projection of miNs tagged with mt-mKeima to identify mitochondria in acidified (left) or neutral (middle) environments. Overlay showing changes in morphology on the right from region seen in white box. **(G)** Quantification of acidified to non-acidified mitochondria in young and old donor miNs treated with DMSO or G2-115. Scale bars 20 µm in A and F. Number of donor lines indicated for each condition, ≥6 neurites collected from ≥3 independent wells per donor line, ≥2 neuronal inductions per donor. Mean ± SEM; analysis by 2-way ANOVA; **p<0.01, ***p<0.001, ****p<0.0001. Graph comparisons which did not meet significance were not marked except within treatment conditions.

To permit characterization of mitochondrial distribution and behavior, mitochondria were imaged in live cells using Mito-Tracker (**Figure S5B**) in combination with LysoTracker Green DND-26 (LysoTracker) which accumulates in acidic compartments to identify autolysosomes (**Video V2**). Organelles in which MitoTracker and LysoTracker co-localized and co-transported (**Video V5**) were categorized as mitophagosomes on resulting kymographs (**Figure 5B**). Treatment with G2-115 or TFEB SA significantly increased the percent of autophagosomes containing mitochondria present in old donor neurons compared to young donor neurons (**Figure 5C**). This implied that when sufficient autolysosomes were present, neurons are correctly identifying damaged mitochondria and targeting them for degradation. Neurons from young donors did not experience a change in mitophagosome density with G2-115 or TFEB SA treatment, presumably because their mitochondria were healthy and functioning appropriately.

Next, we assessed the phenotype of mitochondrial fission and fusion, to determine if improving autophagy and mitophagy changed the dynamics of mitochondria. We found that both mitochondrial fission and fusion events in miNs from old donors were reduced with G2-115 and TFEB SA (**Figure 5D**), improving them to the level seen in healthy young control individuals. G2-115 or TFEB SA had minimal effect on dynamic events in younger donors. To directly evaluate mitophagy, we used the pH sensitive dye mt-mKeima, which undergoes a shift in the wavelength of light required for maximal excitation when mitochondria are in a neutral vs acidic pH (**Figure 5E**). As expected, mitochondria located in an acidified environment (autolysosomes) were small and rounded in shape, while mitochondria in the neutral environment of the cytoplasm were typically elongated (**Figure 5F**). Quantification of the ratio of mitochondria located in an acidified to non-acidified environment showed a significant increase in old donor neurons treated with G2-115 compared to DMSO, consistent with an increase in mitophagy in response to G2-115 treatment (**Figure 5G**). These findings highlight the critical importance of preserving autophagic flux during neuronal aging, and its function as a fulcrum in controlling the behavior of other age-sensitive organelles including mitochondria. Treatment with G2-115 did not significantly alter mitochondrial characteristics such as percent motility, length, or density (**Figure S5C-E**).

To address whether directly modulating mitochondrial fission or fusion would rescue the age-related loss of mitochondrial membrane potential, we exposed miNs from donors of different ages to chemical inhibitors of fission or fusion. We observed that inhibiting mitochondrial fusion by blocking mitofusin-1 using the small molecule MFI8 resulted in high levels of neuronal death after 24 hours of treatment, and complete neuronal loss after 3 days (**Figure S5F**). In contrast, neurons tolerated inhibition of mitochondrial fission using the Drp1 blocker Mdivi-1 for 5 days (**Figure S5F**). Treatment with Mdivi-1 improved mitochondrial membrane potential in old donor miNs compared to untreated old miNs but did not show any meaningful changes in younger cells (**Figure S5G**). The addition of G2-115 to cells treated with Mdivi-1 did not further enhance the membrane potential. Unsurprisingly, mitochondria in miNs treated with Mdivi-1 were substantially longer than those observed in untreated cells (**Figure S5H**) and resulted in a nearly continuous mitochondrial network. Through use of selective inhibitors of mitochondria fission or fusion we identified that blocking mitochondrial fusion is highly detrimental to neurons and results in cellular death, while blocking mitochondrial fission results in an elongated mitochondrial network which acts to temporize the MMP changes observed in fragmented mitochondria during aging.

Finally, we wanted to evaluate the importance of selectively preserving mitophagy against the background of diminished autophagosome acidification during aging. To determine if enhancing mitophagy was sufficient to rejuvenate the behavior and function of mitochondria, miNs from donors of different ages were incubated with the mitophagy induced Urolithin A (UA), gut-microbial metabolite of ellagic acid (Andreux *et al*., 2019; Ryu *et al*., 2016). UA was introduced to cellular media at PID 14 and added with subsequent media changes until imaging. Mature miNs treated with UA showed increased colocalization of MitoTracker and LysoTracker (**Figure S6A**). Quantification of this overlap showed significantly more autophagosomes containing mitochondria in UA-treated cells than DMSO-treated cells, consistent with an increase in mitophagy (**Figure S6B**). Kymographs of tandem-LC3 autophagosomes from miNs treated with UA showed an increase in acidified autophagosomes (**Figure S6C**). The overall percent of acidified autophagosomes did not change in response to UA treatment compared to DMSO treatment in neonatal or young miN lines, but selectively improved in old miNs. However, this improvement was not to the same level observed following total autophagy induction by G2-115 or TFEB SA, and was more donor-dependent than other treatments (**Figure 3J** vs **Figure S6D**). Evaluation of mitochondrial fission and fusion showed a recovery of these dynamic events in response to UA treatment (**Figure S6E**), highlighting that selective mitophagy had a stronger effect on mitochondrial populations than autophagosome populations. Similarly, staining with JC-1 revealed improved mitochondrial membrane potential following UA treatment (**Figure S6F**) in old but not young donor miNs compared to DMSO (**Figure S6G**). Together, these results support evidence that in aged neurons, the increased mitochondrial fission and fusion dynamics reflected the cells need for an organelle-intrinsic mechanism to protect mitochondrial health. The additional feature of decreased autophagosome acidification with aging and simultaneous rescue of autolysosomes and mitochondrial dynamics by increasing autophagy highlighted autophagosome dysfunction as a specific age-related neuronal weakness.

## Discussion

This study presents new insights into dynamic analysis of cellular organelle behavior in live human neurons across an 80-year age spectrum. Leveraging a microRNA-based direct neuronal conversion technique that preserves donor age characteristics, we uncovered age-associated shifts in key organelle pathways. RNA sequencing revealed a downregulation of lysosomal and mitochondrial genes, coupled with an upregulation of genes involved in autophagosome biogenesis, and intracellular trafficking. This correlates with prior work which identified regulatory changes in epigenetics and expression of genes involved in homeostasis of autophagosomes and lysosomes during aging as reflected in the miN system (Chou *et al*., 2025; Lee *et al*., 2024; Oh, Lee and Yoo, 2023; Varghese *et al*., 2025; Lapierre *et al*., 2015). However, the relationship between mitochondrial regulation and genome stability remains less clearly defined, as mitochondrial dysfunction and oxidative stress may promote genome destabilization, potentially establishing a deleterious feedback loop (Feng and Lu, 2025). To validate these transcriptional changes, we employed high-resolution confocal microscopy, which confirmed altered autophagic flux with aging. Remarkably, these deficits were reversible with enhancing autophagic flux in aged neurons. However, aging caused depletion of lysosomes from neurites that remained resistant to pharmacologic correction. Concurrently, mitochondria exhibited age-associated functional alterations. In response, neurons initiated a compensatory upregulation of mitochondrial fission and fusion genes to maintain mitochondrial morphology and spatial distribution. This regulatory change preserved mitochondrial size and distribution within the aging neuron despite the presumed decrease in functional mitophagy resulting from the loss of neuritic autolysosomes (**Figure 6**). Enhancing TFEB-driven processes including autophagy normalized mitochondrial fission and fusion dynamics to that observed in young donor neurons and improved mitochondrial membrane potential. Selectively improving mitophagy in aged miNs likewise improved mitochondrial health and behavior but had a lesser effect on autophagosomes.

**Figure 6:**
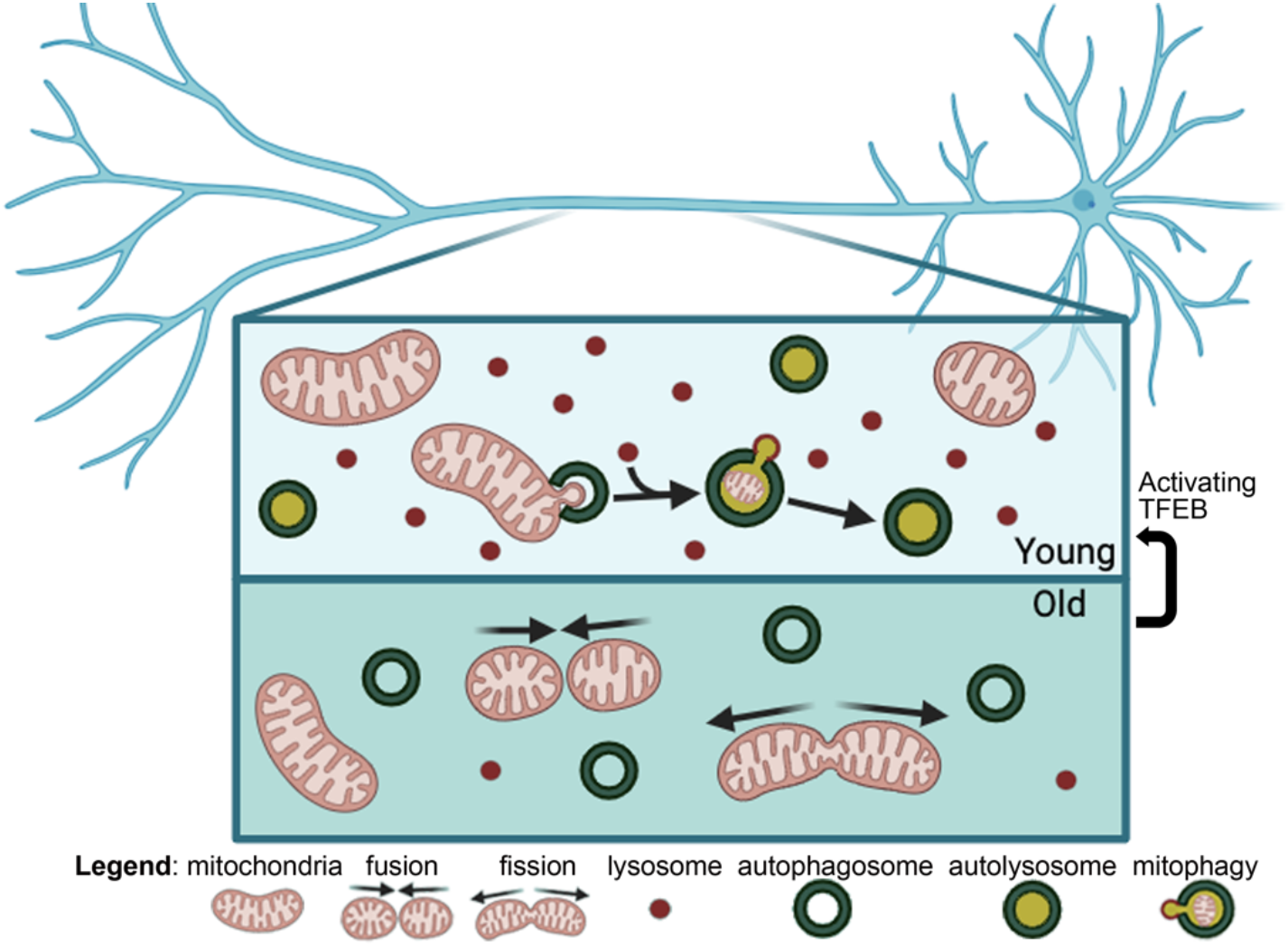
Schematic of autolysosome formation and function in old vs young neurons. TOP: The neurites of neurons derived from young individuals contain many lysosomes (red dots), which can fuse with autophagosomes (green circles) to create functional autolysosomes (yellow). This allows for breakdown and turnover of cytoplasmic components including mitochondria (pink) through mitophagy. BOTTOM: In contrast, the neurites of neurons from old individuals are lacking in sufficient lysosomes, resulting in a deficit of autolysosomes. To preserve their health and integrity, mitochondria increase exchange of membrane and intra-membrane components through fission and fusion in this aged condition.

Incorporation of longitudinally collected isogenic lines confirmed that changes in autolysosome formation and mitochondrial dynamics becomes apparent sometime 65 years of age, although we were not able to pinpoint the timing of this shift. This finding links possible poor neuronal adaptive aging responses in organelles to the age-associated onset of sporadic neurodegenerative conditions. The age at which autophagosome function and mitochondrial self-recovery deviates slightly precedes the average age of onset of Alzheimer and Parkinson diseases (Liang *et al*., 2021). In particular, the observations reported in this work strengthen the hypothesis that mitochondrial dysregulation underlies neuronal fragility and eventual degeneration during unhealthy cellular aging and that this change is required as a byproduct of the failing autophagy pathway.

These findings delineate the processes underlying healthy aging, yet in doing so they highlight areas of neuronal vulnerability. Identification of age-dependent changes in neuronal organelle function and behavior confirms weakness which may confer disease susceptibility in the human brain. Future research directions should address escalating changes associated with disease onset or progression, potentially those downstream of modulation by TFEB including autophagy, trafficking, and fusion, as well as identifying which lysosomal processes are affected. In our system, modulation of TFEB using G2-115 led to a modest effect on lysosomal localization despite TFEB’s role as the master regulator of lysosomal biogenesis via the CLEAR (Coordinated Lysosomal Expression and Regulation) network (Palmieri *et al*., 2011). It is possible that changes within lysosomes occurred, such as an increase in enzyme production, which were not apparent in this study which only evaluated localized expression of LAMP1-positive in neurites, or which we were unable to capture in fixed cells but would be apparent in living cells. Alternately, G2-115 may selectively target a subset of TFEB-based regulatory changes focusing on autophagosomes rather than lysosomes. Clarifying this selective change in autophagosomes but not lysosomes using G2-115 should be further evaluated. Recent work demonstrated that miNs endogenously capture phenotypic hallmarks of Alzheimer disease (Sun *et al*., 2024), further highlighting the power of potential Alzheimer-specific observations associated with aging in future experiments. Identifying organelle changes in this model may prove key to understanding the pathogenesis of this disease in humans, and provide a model for testing early-stage investigative therapeutics.

Our unique method allowed us to elucidate an age-specific decline in human neuronal autophagic flux, and the potential for chemical rescue of this process. We were also able to demonstrate the protective role of mitochondrial fission and fusion as a function of neuronal aging in humans, confirmed by both live cellular imaging and RNA sequencing. We have established a baseline for neuronal mitochondrial length and density in living, non-fixed human neurons, and evaluated their response to aging, although our experimental design was unable to reliably differentiate between axons and dendrites, and unable to determine the distance from the soma due to the requirements of live-cell imaging. Ideally, these findings would be confirmed using 3D reconstruction of fixed brain tissue, although as paraformaldehyde causes mitochondrial fragmentation (Kim *et al*., 2022) this may prove difficult to validate with typical whole-brain fixation methods. This work would also benefit from evaluation of organelle health and behavior in a model including support cells such as glia. Our work highlights that humans may not benefit from selectively enhancing one aspect of mitochondrial dynamics – promoting fission or fusion – to delay aging, but may be amenable to modulation of an upstream regulator such as autophagy or mitophagy to affect larger scale improvement in brain health.

Finally, this work confirms the important neuroprotective role played by autophagosomes, and alternative non-degradative pathways that neurons explore during healthy aging. Promoting autophagy by activating the TFEB system using the small molecule G2-115 or TFEB SA improved neuronal function in otherwise aged healthy neurons, as did improving mitophagy with UA, an exciting finding that suggests that the consequences of aging on neuronal autophagy can potentially be reversed with therapeutics. This result strengthens the general hypothesis that inducing autophagy has the potential to promote healthy aging: one of the frontrunner anti-aging medications, rapamycin (sirolimus), also stimulates autophagy and has previously been demonstrated to extend lifespan in mice (Harrison *et al*., 2009; Juricic *et al*., 2022). Rapamycin is currently used off-label in an effort to slow aging in humans. Further investigation into other pro-autophagy chemicals may identify small molecules capable of protecting neuronal tissue during aging and potentially extending healthy life with fewer off-target effects.

## Methods and Materials

### Fibroblast line and cell culture

Fibroblast lines (Table 1) were purchased or donated (in the case of DIAN (Karch *et al*., 2018)) to the laboratory for use in research. During the fibroblast stage, cells were maintained in DMEM (ThermoFisher Scientific, 11960-044) with 10% or 15% fetal bovine serum (FBS; Gibco, A52567-01) as needed; slower growing cells were given 15% FBS while those that divided quickly were kept in media with 10% FBS. All fibroblast media was additionally supplemented with 1% MEM nonessential amino acids (Corning, 25-025-CI), 1% sodium pyruvate (Corning, 25-000-CI), 1% GlutaMAX (Gibco, 35050-061), 1% 1 M HEPES buffer solution (Corning, 25-060-CI), 1% penicillin/streptomycin (Gibco, 15140-122), and 0.01% beta-mercaptoethanol (Gibco, 21985-023).

### Lentivirus production

Lentivirus used to create cortical miNs required preparation of 5 separate viruses as described below. Creation of each lentivirus-containing plasmid was carried out separately, although all 5 plasmids were transduced at the start of the neuronal conversion process. Production of lentivirus followed previously established protocols (Church *et al*., 2021). In brief, Lenti-X 293T cells (Clonetech) were incubated for 16 hours with polyethyleneimine (PolyScience, 24765-100), the selected plasmid, pMD2.G (envelope), and psPAX2 (empty backbone). Media was exchanged for fresh media (DMEM with 10% FBS, as above) after 16 hours. Supernatant was collected after 72 hours and subsequently kept chilled. Virus-containing supernatant was centrifuged at 1200g for 5 minutes, filtered through a 0.45 µm PES filter membrane (Millex, SLHPR33RB), and centrifuged again at 70,000g for 2 hours. Liquid was aspirated, and the brown viral pellet was resuspended in 1/10^th^ volume cold DBPS, allocated into 1ml tubes, and frozen at -80^°^ C until time of use. Once thawed, virus was only re-frozen once, if necessary, before being discarded.

### Direct neuronal reprogramming

Reprogramming of human fibroblasts to cortical miNs was performed as previously described (Church *et al*., 2021) with few modifications. Briefly, fibroblasts were seeded onto 6-well culture plates (GenClone Cell Culture, 25-105) at a density of 400,000 cells per well. Cells were allowed to attach for 24 hours, then transduced with 5 lentiviruses: miR-9/9*-124, rtTA, p53DD, MYT1L, and NEUROD2 in the presence of 8ug/ml polybrene (Sigma-Aldrich, H9268). 4ml of media containing the viral cocktail and polybrene was added to each well of the 6-well plates, which were then spinfected at 1,000g at 37^°^C for 30 minutes. The following morning, PID 1, cells were rinsed with DPBS and media was exchanged to fresh 10% FBS-containing DMEM with added 1ug/ml doxycycline hyclate (Dox, Sigma-Aldrich, D9891) to induce expression of the microRNA and rtTA. On PID 3, media was again refreshed with the addition of Dox and puromycin dihydrochloride (3 ug/ml, Life Technologies, A11138-03), as well as PID 5 with the further addition of G418 (geneticin sulfate, Life Technologies, 10131-035). At PID 7 cells were replated onto coverslips which had been previously treated with nitric acid, and were prepared with poly-ornithine (Sigma-Aldrich, P4957) and laminin (Sigma-Aldrich, L2020) with fibronectin (Sigma-Aldrich, F4759). On the day of replating, cells were provided fibroblast media with Dox and no antibiotics were given. On PID 8, nascent miNs were changed from fibroblast media to neuronal media (ScienCell Research Laboratories, 1521) supplemented with provided penicillin with streptomycin and growth factor. Neuronal media was made more hospitable for neurons with the addition of 1 μg/mL doxycycline, 200 μM dibutyl cyclic AMP (Sigma-Aldrich, D0627), 1 mM valproic acid, (Millipore, 676380), 200 nM ascorbic acid, (Sigma-Aldrich, A8960), 10 ng/mL BDNF (PeproTech, 450-02), 10 ng/mL NT-3 (PeproTech, 450-03), 1 μM retinoic acid (Sigma-Aldrich, R2625), 100x RevitaCell (Life Technologies, A2644501), and 3 μg/mL puromycin. A 50% media change was performed on PID 10 and 14, and an additional treatment with Dox was provided on PID 12. After PID 14 miNs were considered mature enough to discontinue selection with Dox and puromycin; RevitaCell was also stopped at this time. Additional 50% media changes were performed every 4 days until cells were mature at PID 30. All imaging was done between PID 28 and PID 32 to balance the constraints of cellular maturity against the time required for live-cell imaging.

### mt-mKeima treatment

The pHAGE-mt-mKeima was obtained from Addgene (#131626) and used for lentiviral transduction. To ensure the purity of the virus product and higher concentration of virus, supernatant collected from 293T cells was filtered as normal through a 0.45 μm PES membrane and incubated overnight incubation with Lenti-X concentrator (TakaraBio 631231). The resulting slurry was centrifuged at 1,500 g for 45 minutes at 4o and resuspended in 1/10^th^ of the original volume with PBS. The 10x concentrated virus was then gently floated on 7 ml of a 20% sucrose cushion (20% sucrose, 100 mM NaCl, 20 mM HEPES [pH 7.4], 1 mM EDTA in distilled water) in an ultracentrifuge tube and centrifuged at 70,000 g for 2 hours at 4^°^ C to pellet the virus (Oh, Lee and Yoo, 2023). The resulting double-purified virus was resuspended in 1/100^th^ of the starting volume in cold PBS. For each well of a 96 well plate, 0.5 μl of the virus preparation was added to the culture media 6 days prior to imaging. Media changes proceeded as normal.

### Tandem LC3 treatment

To express tandem mCherry-GFP-LC3 in miNs, ultra-concentrated virus containing the tandem construct (a gift from Anne Brunet) was added to culture media 6 days prior to imaging (PID 24) as previously described (Lee *et al*., 2024). Media changes proceeded as normal thereafter.

### Autophagy induction with G2-115

Induction of autophagy was conducted using the small molecule G2-115, a kind gift from David Perlmutter at Washington University. Steps in synthesis of G2-115 are briefly summarized in a recent publication (Oh *et al*., 2022). The addition of G2-115 followed previously established protocols (Lee *et al*., 2024; Oh *et al*., 2022). In brief, 1uM G2-115 in DMSO was added to the developing miNs on PID 15 during a 50% media change to reach a final concentration of 0.5uM. Thereafter, each media change included 0.5uM G2-115 through the end of culture at PID 30. As DMSO has been shown to alter the behavior of cultured cells in high concentrations, equal concentration DMSO was added to control wells in parallel to G2-115.

### Autophagy induction with TFEB SA

The TFEB SA construct, which contains two mutated residues S142 and 211A has been previously described (Lee *et al*., 2024). Briefly, TFEB wildtype pcDNA3.1-TFEB-WT-MYC (#99955) was obtained from Addgene, mutagenized (S142/211A), and ligated into the N174-lentiviral vector. Virus production was performed as described above, and double-concentrated TFEB SA was added to miNs on PID 14. Media changes proceeded as normal thereafter.

### Mitochondrial fission and fusion modulation

MFI8 (MedChemExpress HY-150031) and Mdivi-1 (MedChemExpress HY-15886) were resuspended in DMSO per manufacturer’s instructions and stored at -80^°^ C until use. Active aliquots were stored at -20^°^ C for no longer than one week. MFI8 was tested for 1, 3, 5, and 7 days and found to kill cells within 24-72 hours at a concentration of 20 μM, a dose commonly used in the literature (Zacharioudakis *et al*., 2022). Notably, prior work with MFI8 had recommended treatment for 6 hours, which would have been insufficient to counteract the two-week long treatment period for G2-115. Mdivi-1 was likewise tested at a concentration of 10 μM in line with prior studies (Cui *et al*., 2010; Gan *et al*., 2014; Xu *et al*., 2016) for 1, 3, 5, and 7 days, with fresh chemical added every 48 hours due to its short half-life. Cells survived through day 7 but appeared stunted; as such 5 days of treatment was chosen for further experiments.

### Mitophagy induction with Urolithin A

Urolithin A (MedChemExpress HY-100599) was resuspended in in DMSO per manufacturer’s instructions and stored at -80^°^ C until use. Active aliquots were stored at -20^°^ C for no longer than two weeks. Urolithin A was introduced on PID 14 to a final concentration of 5 μM in the well (2x added with the first media change) and fresh Urolithin A was added with each media change (1x) thereafter. A concentration of 5 μM was chosen as it aligned with previous studies in neuronal-type cells (Kim, Lee and Kim, 2020; Zhang *et al*., 2025).

### Immunocytochemistry

Cells were fixed with 4% paraformaldehyde (Electron Microscopy Sciences, 15710) for 20 minutes at room temperature and then rinsed 3 times with PBS. For coverslips not needed immediately, cells were immersed in PBS, wrapped in parafilm, and stored at 4^°^C. To stain, cells were permeabilized and blocked for 1 hour in PBS containing 5% BSA (Sigma-Aldrich, 0000484062), 0.3% Triton X-100 (Sigma-Aldrich, T8787), and 2% goat serum (Sigma-Aldrich, G9023) and then incubated overnight at 4^°^C in block solution containing primary antibody. The next day, fixed cells were washed three times with wash buffer containing 5% BSA with 0.3% triton X-100 and then incubated for 1 hour at room temperature with secondary antibody in wash buffer. Stained cells were rinsed once with PBS and then labelled with DAPI for 5 minutes. Cells were rinsed a final time with PBS and then mounted onto coverslips using ProLong Gold antifade (Invitrogen, P36934) mounting agent.

Primary antibodies used for immunostaining include rabbit anti-Tau (Agilent/DAKO, A002401-2 1:200), chicken anti-TUBB3 (TUJ1, Novus, NB100-1612 1:1000), mouse anti-LC3 (Cell Signaling, 83506; 1:100), rabbit anti-TFEB (Cell Signaling, 4240, 1:750), and rabbit anti-LAMP1 (Abcam, ab24170; 1:200). Secondary antibodies were IgG raised in goat and conjugated to fluorophores (Invitrogen). This included anti-mouse, anti-rabbit, and anti-chicken conjugated to Alexa-488, Alexa-568, and Alexa-647.

### Fixed-cell confocal microscopy and analysis

Images of immunostained cells were captured using a Leica S5X white light laser confocal system with Leica Application Suite (LAS) Advanced Fluorescence 2.7.3.9723. For each coverslip, three images in separate fields of view were collected per sample at 100x magnification for lysosomal quantification. Images were collected as a z-stack with a spacing of 0.4 µm and projected in FIJI to allow collection of maximal fluorescent data. Lysosome analysis was performed using open-source FIJI software. Two neurites were selected per field of view. A region of interest 20 pixels wide (approximately 10 pixels to each side of the neurite) tracking each neurite was collected and stored for both the lysosome channel (LAMP1 antibody) and the tubulin channel (TUJ antibody). Neurite images were thresholded in FIJI using the Huang method, which allowed for detailed inclusion of the neurite using the Analyze Particles features. The identified regions of interest (tubulin-positive neurite) were then applied to the LAMP1 or LC3 and analyzed for signal intensity. To calculate overall intensity, a weighted average by neurite area was calculated per image, then averaged across three images per cell line. The JACoP co-localization plugin was used to calculate the Mander’s M1 coefficient between LAMP1 and TUJ signals. A total of 6 neurites of data were collected from each donor cell line, with three cell lines per condition as available. Results from the neurites were averaged over one cell line; further calculations and statistics were performed in Prism (as below). Visualization of tau staining to determine maturity of cortical miNs was also performed using FIJI. TFEB signal was captured with a 40x objective; images were taken from five or more fields of view. Nuclear DAPI signal was used for thresholding in FIJI, then regions of interest (nuclei) were selected using Analyze Particles and applied to the TFEB channel. Intensity of each nuclei was recorded and background subtracted for 488 signal in the field. Over 100 nuclei per condition were recorded.

### Live cell confocal

Live cell imaging was performed on a Zeiss CellDiscoverer 7 equipped with an LSM900 scan head and two Colibri LED lasers as well as two GaAsP PMTs for simultaneous color detection. The microscope additionally provides a sealed heat-, CO_2_-, and humidity-controlled chamber for prolonged imaging without compromising cell health. For each cell line, at least two wells from at least two separate reprogramming inductions were used. Two videos (MitoTracker/LysoTracker and tandem-LC3) or 2-3 images (JC-1 and mt-mKeima) were captured for each well of a glass-bottomed 96-well plate. The data from each well was averaged for analysis, and the pooled averages were used in the final calculations for each cell line. All videos were captured with dual illumination in red and green channels simultaneously at 50x magnification. Videos were 5 minutes long, with one frame taken every three seconds. Details on still-images for JC-1 and mt-mKeima are below.

For recordings of mitochondria and mitophagosomes, mature cortical miNs were incubated with 200nM MitoTracker™ Red CMXRos (Invitrogen, M7512) and 100nM LysoTracker™ Green DND-26 (Invitrogen, L7526) for 30 minutes. Fresh media was added, and cells were left for 30 minutes to equilibrate. Stained cells were then imaged on the CellDiscoverer 7. For each video (**Video V2**), analysis was performed on two neurites, which were used to generate kymographs in both the red and green channels as previously established (Klinman and Holzbaur, 2016). Selected neurites met criteria of not having neither visible associated cell bodies nor neurite terminals within the field of view (80.27×80.27 μm), appearing mature (thin) and straight, and if possible, having at least one motile mitochondrion to improve ease of tracing the span of the chosen neurite. The kymographs were used to determine the length of each mitochondrion, the percent of motile mitochondria, number of autophagosomes per neurite, number of autophagosomes which contained mitochondrial signal, and the length of the neurite (**Video V2**). Values for fission and fusion were separately calculated for all visible neurites in each field of view, not per kymograph (**Video V3**). Each event was determined if two mitochondria came into contact or separated for more than 6 seconds (two video frames). Values for total number of fission or fusion events were divided by the total number of starting mitochondria in the field of view, allowing calculation of percent of mitochondria undergoing fission or fusion during the video. These values were then averaged to provide total percent of fission or fusion events. As with other mitochondrial calculations based on kymographs, averages for fission and fusion were calculated on a per-well basis and then further averaged with other wells from the same cell line. Results were reported as percent total events per distinct mitochondria present on the first frame of the recording, as the raw number of neurites and thus mitochondria varied between recording locations.

Live-cell recording using the tandem mCherry-GFP-LC3 construct was performed in a similar fashion. Mature cortical miNs that had been transduced with lentivirus containing the tandem LC3 construct 6 days prior were imaged alive in the CellDiscover 7 on PID 30. Resulting videos, again two per well of a 96 well plate, three or more wells imaged per cell line, were analyzed for movement, density, and acidification of the fluorescently labeled autophagosomes following selection of neurites which met the same criteria as used in MitoTracker selection (**Video V4**). Quantification of autophagosome properties was performed using only the red channel, as this captured both acidified and non-acidified organelles. Kymographs generated from two neurites per field of view in both the red and green channel were used to determine the percent of identified autophagosomes (red) which were not acidified (green). Results were averaged on a per-well basis and then over the number of sampled wells for each cell line.

Evaluation of mitochondrial health using color-shifting fluorescent JC-1 marker (Invitrogen, M34152A) was also performed in live cells. As with MitoTracker, mature cortical miNs were incubated with 2μM JC-1 in neuronal media for 30 minutes on the day of imaging. Cells were washed once with PBS to remove unbound probe, and then placed back in conditioned neuronal media for imaging on the CellDiscoverer 7. Two or three separate fields of view per well were collected as a z-stack with a vertical spacing of 0.3μm using the 50x confocal objective. The z-stacks were compressed into a max intensity projection using FIJI, and the mean intensity of the red and green channels was calculated per image; resulting data was corrected for background intensity. A total of 3 or more independent wells per condition were summed to generate the final fluorescence intensity per donor line, which was analyzed in Prism.

Mitophagy was evaluated using mt-mKeima, which was introduced to the neurons 6 days prior to imaging. On the day of imaging, mature cortical miNs were imaged on the CellDiscover7. Two wells per cell line from two separate inductions were used, for a total of 4 wells. The miNs were first exposed to 405 nm light for the full z-stack (50x objective, 0.3 μm spacing between images) and emission was detected at 620 nm. A second z-stack was then collected with excitation using 594 nm light and emission in the same 620 region. The z-stacks were compressed into a maximum intensity projection in FIJI, and the mean intensity of the 405-excitation and 594-excitation (pseudo-colored cyan and magenta for ease of view) were calculated per image and corrected for background intensity. The averages of each well were summed and imported to Prism for graph generation and analysis.

### Live cell recording of neuronal reprogramming

Recording of the conversion of fibroblasts to miNs over the span of 21 days took place using an IncuCyte S3 LiveCell Analysis System and corresponding IncuCyte software (Sartorius). In brief, fibroblasts were cultured per normal protocols and infected with neuron-promoting virus on PID 0. They received induction with doxycycline on PID 1 and 3, and selection with puromycin on PID 3. Replating took place on PID 4, with cells distributed to a 96-well glass-bottom dish. Cells remained in fibroblast media through PID 6, when media was changed to neuronal media. Feedings were continued per standard schedule after conversion to neuronal media. At time of viral infection (PID 0), an additional well was treated with RFP expressed under the CMV promotor. These RFP-expressing cells were mixed at 5% or 10% with non-fluorescent cells for ease of cell tracking during the course of direct neuronal conversion (**Video V1**).

### RNA sequencing analysis

The results generated through RNAseq relied on re-analysis of a previously collected data set. The initial data was analyzed for differentially expressed genes (DEGs) between healthy old and healthy young donor cells, for details please see reference (Sun *et al*., 2024), Supplemental Figure 12B and Supplemental Table 9. Cortical cells from 4 healthy young and healthy old individuals were grown in 3D culture and collected on PID 30 for RNA analysis. The transcriptome of 3 biological replicates were analyzed for each donor. For details of quality control and sequencing validation please see (Sun *et al*., 2024). Resulting DEGs were re-analyzed to obtain adjusted p value. Genes with adjusted p value <0.01, base mean >100, and |log2fold change| >0.58 were included for further gene ontology (GO) analysis of cellular component and biologic process groupings using PANTHER through GeneOntology (https://geneontology.org/). Duplicate entries (>50% title similarity and >90% gene inclusion similarity) were removed. Hand-selection of specific GO terms involved in organelle behavior which reached statistical significance were included in Figure S2A-B. The 15 ontologies with the largest different between the old and young data sets are included in Figure S2C and D.

### Quantification and statistics

Data generated from fixed-cell imaging and live-cell imaging was stored in Excel files and subsequently imported into GraphPad Prism v10 for quantification and statistical analysis. Statistical comparisons were performed using two-way analysis of variance (ANOVA) or t-test. Fixed cell imaging and analysis of lysosome localization was performed blinded and confirmed by a second independent investigator. Data was collected on the same day across all cell lines whenever possible, and analysis was performed by the same operator on congruent data sets within one week.

Cell lines were chosen based on availability. Six longitudinal male donors were chosen as being the only fibroblast lines available through Coriell spanning 15-16 years apart in time of collection and divided by first collection <35 years old or last collection >80 years old. Only one female longitudinal line met the same criteria; this data is available if requested but was not included in calculations due to lack of additional lines for comparison. Non-isogenic female lines were chosen due to availability within the laboratory and similarity in age to the youngest and oldest of the male samples. Only two neonatal lines were available; although other lines were available for purchase, it was not clear that they had different initial source tissue from the two included lines. No female neonatal lines were available. Sample size was determined based off initial power calculations from mitochondrial fission experiments in healthy young and old donors, assuming statistical power of 0.80 and an alpha of 0.05. At the time, it was predicted that 6 patient samples per condition would be needed. As such, 3 male and 3 female cell lines were used where available. The experiments were not powered to determine differences between donors of different sexes, although some results did indicate significant changes in this dimension.

All graphs are expressed as mean± SEM from at least three independent cell lines unless otherwise noted (only two lines available for neonatal quantification). Differences were considered statistically significant at *p<0.05, **p<0.01, ***p<0.001, ****p<0.0001. Detailed statistical information for each experiment is provided in the corresponding figure legend. Incomplete data sets (less than three independent wells of live cell imaging available for a cell line) were excluded: this resulted in loss of occasional data from one cell line (AG08790) due to the very slow growth of the fibroblasts; as such insufficient cells were available to generate healthy miNs in sufficient quantities to carry out all experiments, despite growing the fibroblast sample for >5 months.

## Acknowledgements

The authors are grateful to Dr. Jerome Molleston and Dr. Courtney Walker, who provided thoughtful suggestions and textual edits to this work. This study was supported by the following programs, grants and fellowships: WashU PMI-Phase II (A.S.Y.), NIH/NIA R01AG078964 (A.S.Y.), Cure Alzheimer’s Fund (A.S.Y.), Longer Life Foundation (E.K.), McKnight Brain Research Foundation through the American Brain Foundation and the American Academy of Neurology (E.K.), Alzheimer’s Association (E.K.) and Research Education Component of NIH/NIA P30AG066444 (E.K.). Data collection and sharing for a fibroblast line used in this project was supported by The Dominantly Inherited Alzheimer Network (DIAN, U19AG032438 & 3U19AG032438-12S2) funded by the National Institute on Aging (NIA), the Alzheimer’s Association (SG-20-690363-DIAN LATAM & DIAN OBS-26-1552363), the German Center for Neurodegenerative Diseases (DZNE), Raul Carrea Institute for Neurological Research (FLENI), Partial support by the Research and Development Grants for Dementia from Japan Agency for Medical Research and Development (AMED), the Korea Health Technology R&D Project through the Korea Health Industry Development Institute (KHIDI), Korea Dementia Research Center (KDRC), funded by the Ministry of Health & Welfare and Ministry of Science and ICT, Republic of Korea RS-2024-00344521, and Spanish Institute of Health Carlos III (ISCIII). This manuscript has been reviewed by DIAN Study investigators for scientific content and consistency of data interpretation with previous DIAN Study publications. We acknowledge the altruism of the participants and their families and contributions of the DIAN research and support staff at each of the participating sites for their contributions to this study.

## Supplemental Figures

**Supplemental Figure S1:**
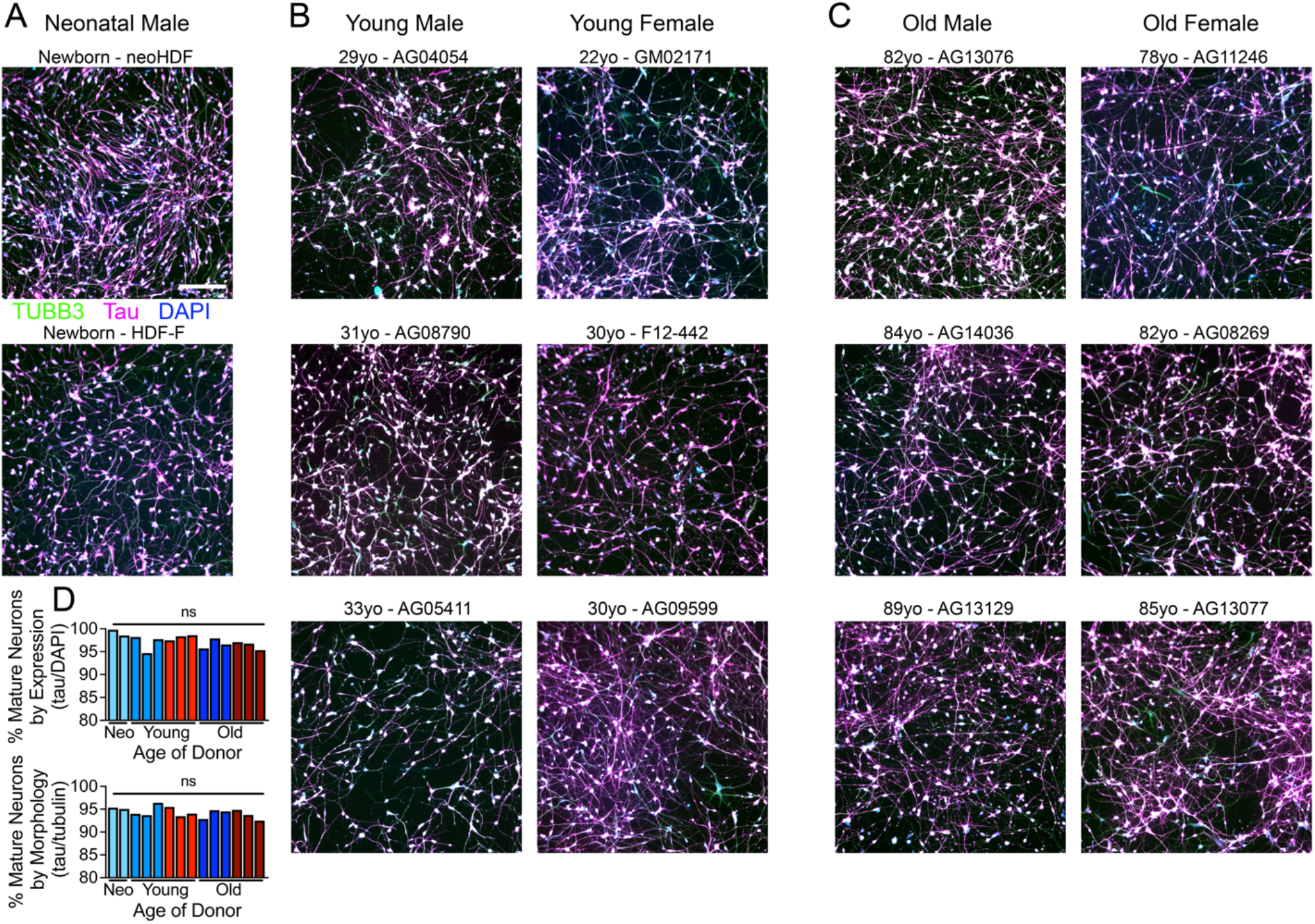
microRNA-induces reprogramming across fibroblast lines independent of donors’ ages or sex. A. Representative confocal images of two reprogrammed neonatal lines used throughout the manuscript, referred to as neoHDF and HDF-F, stained with TUBB3 (tubulin, morphology marker), tau (neuronal marker), and DAPI. B. Representative confocal images from young male and young female reprogrammed miNs, donor ages 22 through 33 years old, stained for TUBB3, tau, and DAPI. C. Representative reprogrammed miNs on confocal imaging from old male and female donors aged 78 through 89 years old, stained for TUBB3, tau, and DAPI. D. Quantification of reprogramming efficiency by age, calculated by percent of DAPI-positive cells which express tau (top) and by morphology using percent of TUBB3-positive cells which express tau (bottom). Donor line colors correspond to donor ID in Table 1. Scale bar 100µm, same magnification throughout. At least 200 cells per field of view per donor cell line used for quantification of neuronal maturation.

**Supplemental Figure S2:**
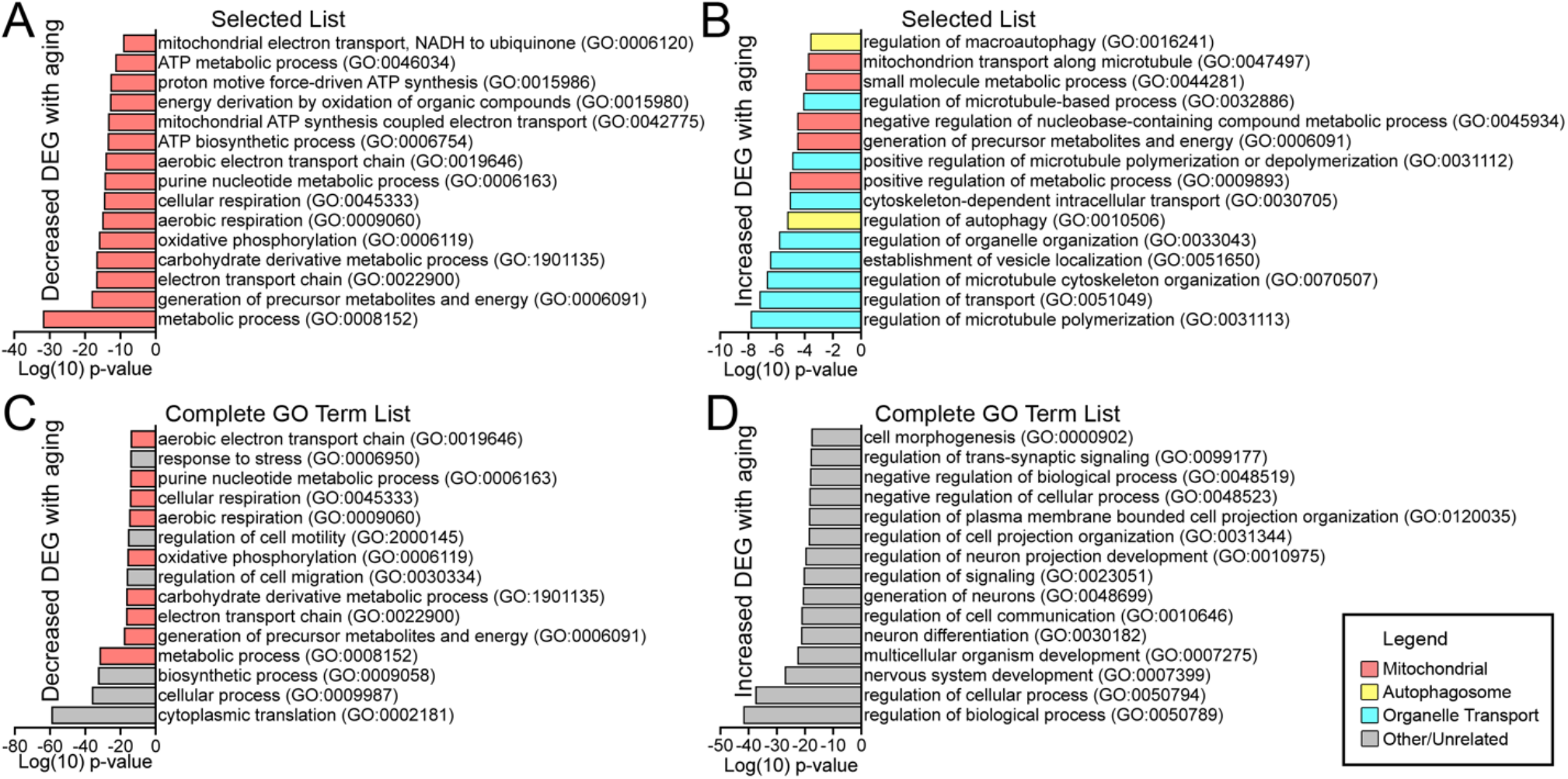
Expression of organelle-associated genes changes with age by RNA sequencing and gene ontology. A. Significance score of differentially expressed genes (DEG) downregulated with aging from RNAseq sorted by gene ontology (GO) biological process terms. B. Significant GO terms related to organelle and transport DEGs as in A which increase transcript expression as a function of aging. C. Top 15 biological processes GO terms which are downregulated with age as determined by PANTHER. D. Top 15 biological process GO terms as in C which are up regulated with age. All included DEGs met criteria for adjusted P < 0.01, |log_2_fold change| > 0.58, and baseMean > 100. Legend below; mitochondrial (pink), autophagosomal (yellow), transport processes (blue), or unrelated (grey).

**Supplemental Figure S3:**
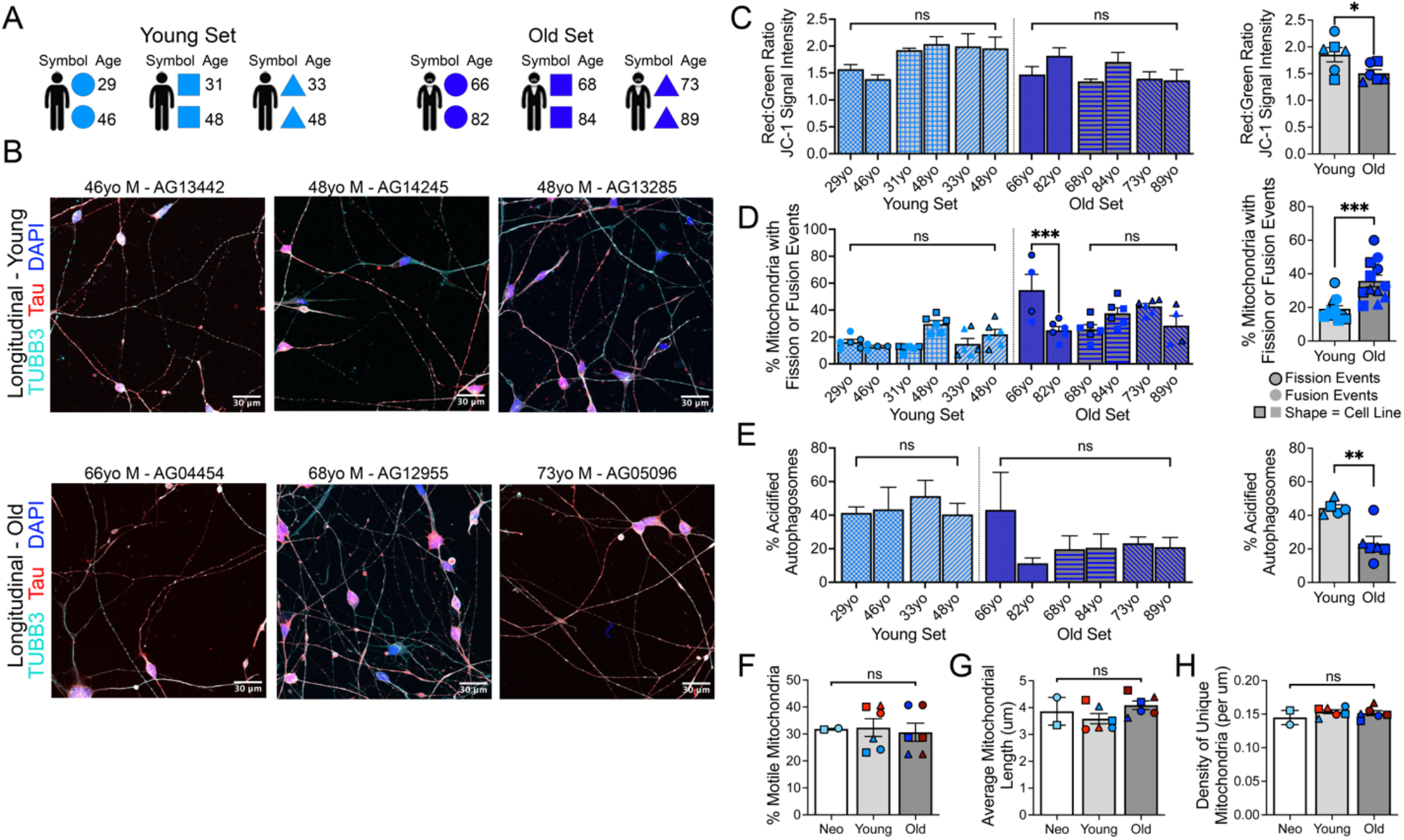
Mitochondrial aging and dynamics in young and old grouped longitudinal samples. A. Schematic of longitudinally collected samples from younger and older individuals collected 15-16 years apart. Younger lines on the left, older on the right. No female lines available with skin biopsy ≥15 years between donations. B. Longitudinally collected male fibroblasts reprogram into robust miNs, stained with tubulin, tau, and DAPI. Scale bars 30 µm. C. Changes in MMP with age in isogenic male samples using JC-1 color-change fluorescent assay. Divided by isogenic donor on the left, collapsed by age group on the right. D. Quantification of dynamic fission and fusion events in longitudinal male samples. Fission depicted as outlined symbols, fusion as symbols without online of the same shape from the same cell line. Divided by isogenic donor on the left, collapsed by age group on the right. E. Quantification of percent autophagosome acidification in isogenic male samples using tandem-LC3 construct. Per donor on the left, collated by age group on the right. F. Percent of of mitochondria which move >5 µm along neurites in 5 minutes in miNs derived from donors of different ages. G. Quantification of length of mitochondria in neurites from miNs derived from donors of different ages. H. Quantification of mitochondrial density in neurites of miNs from donors of different ages. ≥6 neurites collected from ≥2 independent wells per cell line. Six total male donors contributed longitudinal samples, 3 young and 3 old with ages at time of fibroblast donation noted to the right. Mean ± SEM; 2-way ANOVA; *p<0.05, **p<0.01, ***p<0.001

**Supplemental Figure S4:**
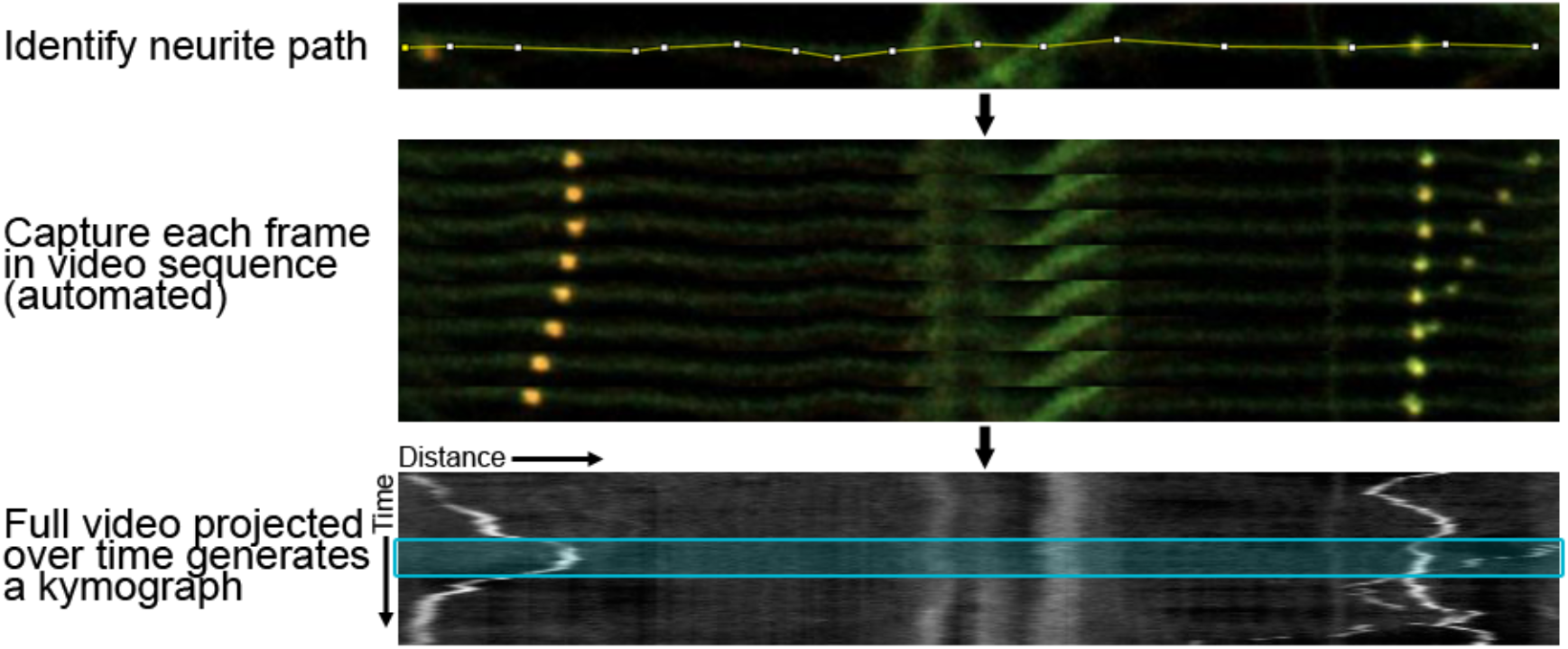
Kymograph generation from video recording of organelle behavior in living miNs derived from patient fibroblasts. (TOP) Path of a single neurite is identified, and carefully traced using segmented line tool in FIJI. (MIDDLE) To create the resulting 2D projection kymograph of movement along the neurite selected, individual frames are stacked on top of each other. The process is automated in FIJI, here 20-pixel segments from 8 sequential still frames are shown to highlight the movement of the three autophagosomes (yellow dots) along the selected neurite. (BOTTOM) The resulting kymograph depicts the position (x-axis) of each fluorescent organelle as a function of time (y-axis), here shown in white-on-black. Three autophagosomes are observed in this example. Aqua box in the middle of the field corresponds to the separated still frames shown in MIDDLE.

**Supplemental Figure S5:**
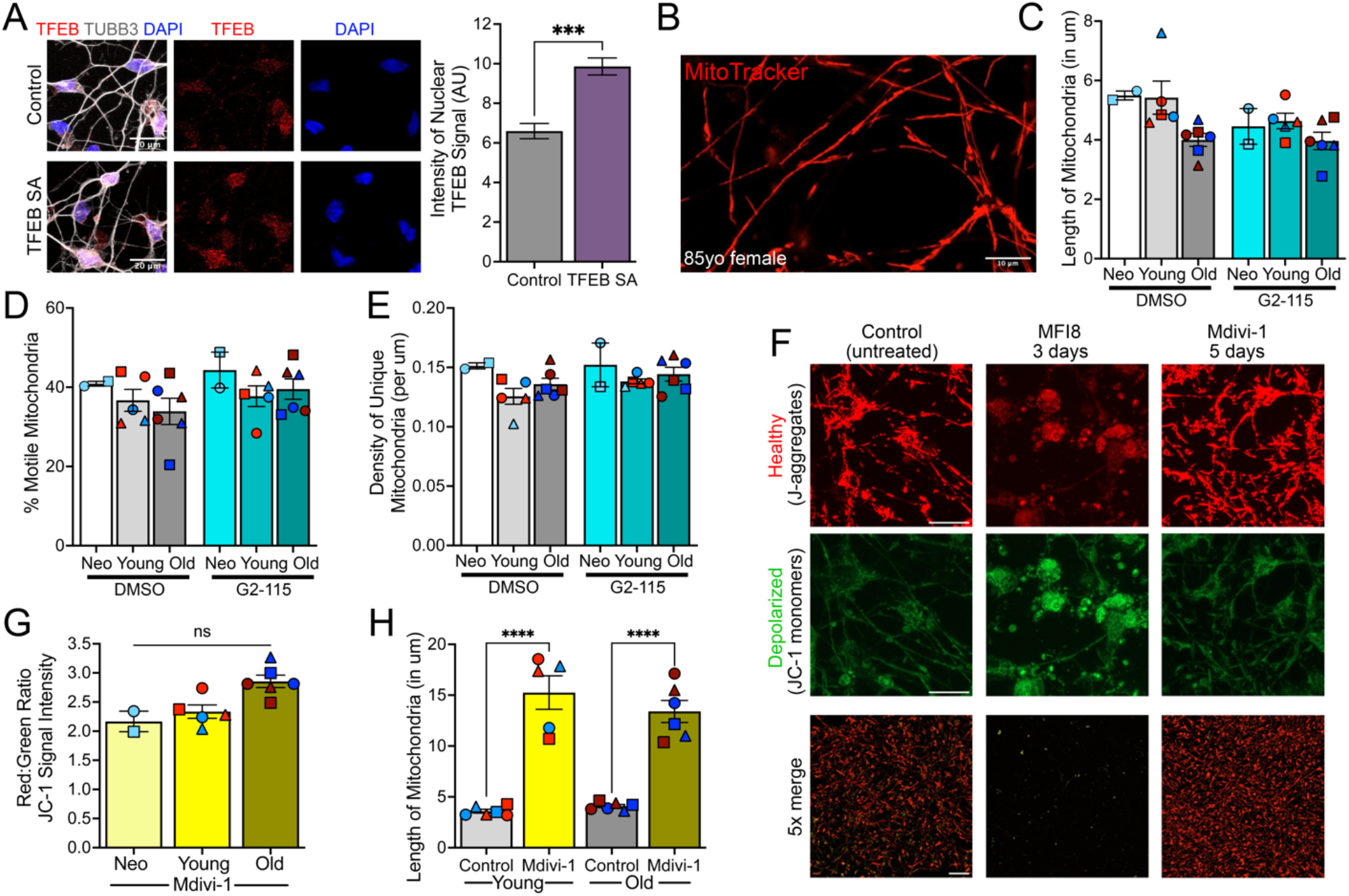
Selected mitochondrial characteristics do not respond to induction of autophagy but do respond to inhibition of mitochondrial fission or fusion. A. Representative images (left) and quantification (right) of nuclear TFEB from cortical control or TFEB SA-trans-duced miNs immunostained with anti-TFEB and TUBB3 antibodies. An average of 100 cells per condition were counted from >5 randomly chosen fields. Scale bars, 20 μm. B. Representative still frame from live-cell recording of MitoTracker (red) labeled mitochondria in miN from an 85-year-old female donor. Scale bar 100 µm. C. Quantification of length of mitochondria in neurites from miNs derived from donors of different ages treated with DMSO or G2-115. D. Percent of mitochondria which move >5 µm in 5 minutes from donors of different ages treated with DMSO or the autophagy inducer G2-115. E. Quantification of mitochondrial density in neurites of miNs treated with G2-115 or DMSO from donors of different ages. F. JC-1 imaging of PID 30 miNs with or without mitochondrial fusion inhibitor MFI8 or mitochondrial fission inhibitor Mdivi-1. 5x resolution imaging shows overall density of surviving cells, bottom row. Scale bar 20 µm top two rows and 100 μm 5x merged images. G. Quantification of JC-1 signal intensity in donor miNs of different ages treated with Mdivi-1 for 5 days. H. Quantification of mitochondrial length calculated using MitoTracker in miNs treated with Mdivi-1 or control. ≥6 neurites collected from ≥3 independent wells per donor line (C-E), miN lines differentiated at least twice per donor (C-H). Mean ± SEM; 2-way ANOVA (C-H), t-test (A); no significant difference detected between cells of different ages or those treated with DMSO and G2-115 in C-E. ***p<0,01, ****p<0.001.

**Figure S6:**
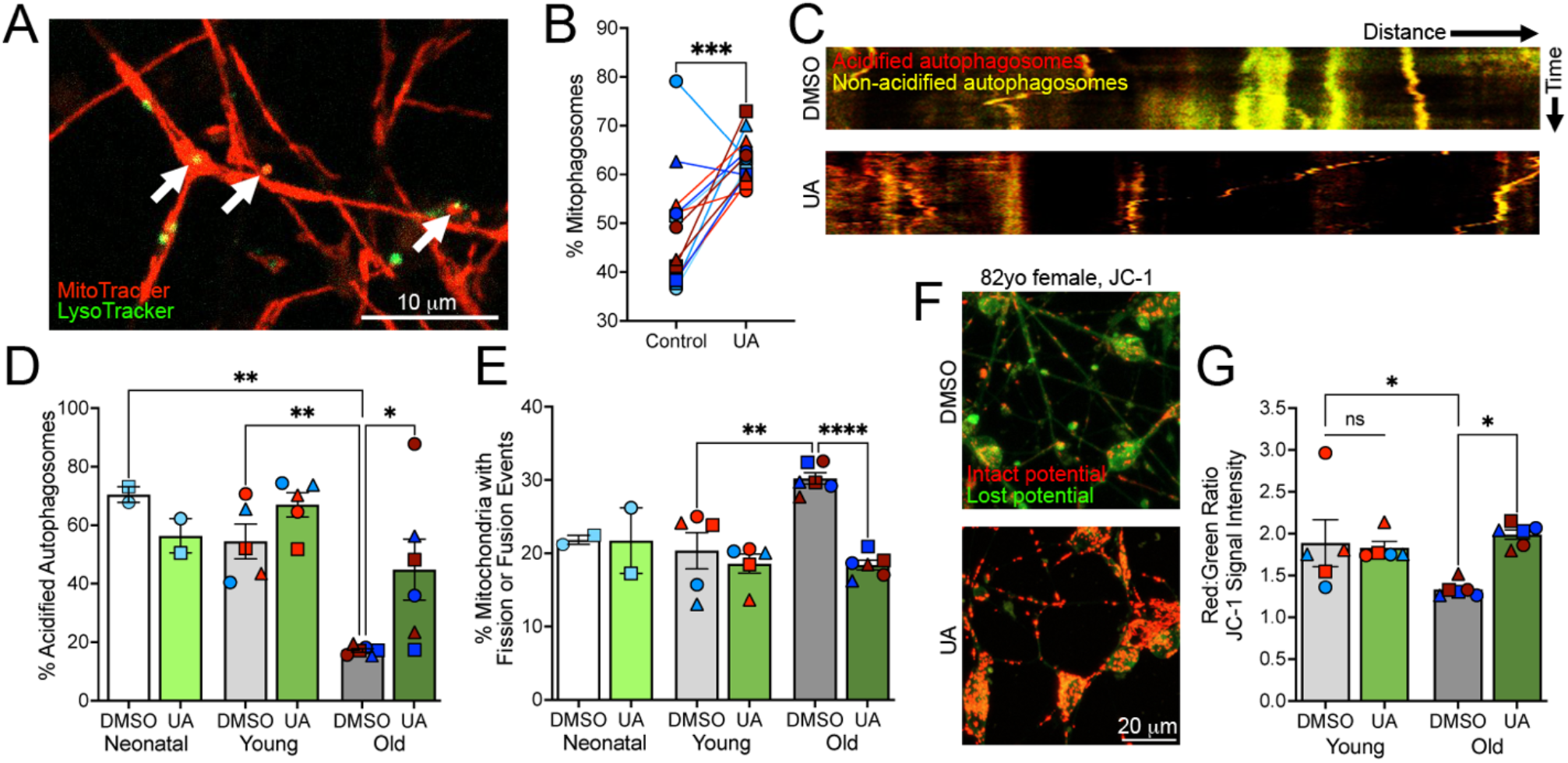
Treatment with Urolithin A increases mitophagy in old donor miNs resulting in recovery of mito-chondrial health. A. Representative still frame from live-cell recording of MitoTracker (red) mitochondria and LysoTracker (green) autophagosomes in miN treated with Urolithin A (UA). White arrows indicate mitophagosome. Scale bar 10 µm. B. Quantification of percent of autophagosomes which contain mitochondria fragments across all donor lines in response to treatment with UA. C. Kymograph of tandem-LC3 behavior in miNs treated with DMSO (top) or UA (bottom); kymographs oriented with cell bodies presumably to the left consistent with retrograde motion of autophagosomes. D. Quantification tandem-LC3 data evaluating percent of acidified autophagosomes in miNs from donors of different ages treated with DMSO or UA. E. Quantification of mitochondria percent fission and fusion using MitoTracker staining (as in A) in miNs from donors of different ages treated with DMSO or UA. F. Z-projection of live mitochondria labeled with JC-1 to identify healthy mitochondria (red) and those with depolarized membranes (green) in miNs treated with DMSO or UA. Scale bar 20 µm. G. Quantification of JC-1 data in young and old donor miNs treated with DMSO or UA. Each donor line was differentiated twice per experiment, ≥4 independent wells per donor were analyzed, and ≥8 neurites collected for B, D, and E. Mean ± SEM; 2-way ANOVA; *p<0.05, **p<0.01, ***p<0.001, ****p<0.0001.

## Video Files – Please contact for files

**Video V1**: Recording of cells undergoing transition from fibroblast to miN, expressing exogenous RFP-tagged synapsin in 5% of total cells. Video begins at time of replating on to glass 96-well dish, PID 4. Recording taken once per hour using IncuCyte LiveCell system. Selected still frames from recording are show in Figure 1B.

**Video V2**: Live-cell recording of representative cortical miN stained with MitoTracker red to label mitochondria (red, left) and LysoTracker green to label acidic organelles (green, right). Recording taken at 1 frame every 3 seconds for 5 minutes. Video recorded from cell line AG09599, 30-year-old female.

**Video V3**: Live-cell recording of mitochondrial dynamics from miN neurites demonstrating an episode of fission (left) and fusion (right) with corresponding kymographs. MitoTracker labelled mitochondria in red undergo fission and fusion over the course of the 5-minute recording (top), which are visible as regions where a single black-colored mitochondria divides into two, or two mitochondria fuse into one in the resulting kymograph (bottom). Fission recording taken from line AG13129 an 89-year-old man, and fusion recording taken from line AG04054 a 29-year-old man.

**Video V4**: Live-cell recordings of individual neurites in miNs from donors of different ages expressing the mCherry-GFP-LC3 tandem construct. Videos recorded at 1 frame every 3 seconds for 5 minutes. miNs from young donors on the left, miNs from old donors on the right. Female donors at the top, male donors on the bottom. Red dots indicate autophagosome, green dots indicate lack of acidification (retained GFP signal). In merged videos (yellow), yellow dots signify unacidified autophagosomes, while red dots signify acidified autolysosomes.

**Video V5**: Live-cell recording of individual neurites in miNs from donors of different ages stained with MitoTracker (red) and LysoTracker (cyan), overlay on bottom of each set highlights mitophagosomes (white). Recordings on the left from male donors, and on the right from female donors. Donor age advances from top (neonatal) to bottom (age

>75 years) within each column. Top left recording from line HDF-F, neonatal male; middle left recording from line AG05411, 33-year-old male; bottom left recording from line AG13076; 82-year-old male; top right recording from line AG09599, 30-year-old female; bottom right from line AG13077, 85-year-old female.

## Notes

### Competing Interest Statement

The authors have declared no competing interest.

### Summary of Updates

Addition of mechanism to explain the function of drug G2-115 including the role of TFEB. Experiments containing obligate-nuclear TFEB added. Experiments involving pH of mitochondrial environment added. Extensive additional drug compound testing added including Urolithin A, mitochondrial fission inhibitor, and mitochondrial fusion inhibitor. Discussion and results sections updated to reflect new data and new analysis of existing data (lysosomes).

## References

Abernathy, D. G., Kim, W. K., McCoy, M. J., Lake, A. M., Ouwenga, R., Lee, S. W., Xing, X., Li, D., Lee, H. J., Heuckeroth, R. O., Dougherty, J. D., Wang, T. and Yoo, A. S. (2017) ‘MicroRNAs Induce a Permissive Chromatin Environment that Enables Neuronal Subtype-Specific Reprogramming of Adult Human Fibroblasts’, Cell Stem Cell, 21(3), pp. 332–348.e9.

Aman, Y., Schmauck-Medina, T., Hansen, M., Morimoto, R. I., Simon, A. K., Bjedov, I., Palikaras, K., Simonsen, A., Johansen, T., Tavernarakis, N., Rubinsztein, D. C., Partridge, L., Kroemer, G., Labbadia, J. and Fang, E. F. (2021) ‘Autophagy in healthy aging and disease’, Nat Aging, 1(8), pp. 634–650.

Andreux, P. A., Blanco-Bose, W., Ryu, D., Burdet, F., Ibberson, M., Aebischer, P., Auwerx, J., Singh, A. and Rinsch, C. (2019) ‘The mitophagy activator urolithin A is safe and induces a molecular signature of improved mitochondrial and cellular health in humans’, Nat Metab, 1(6), pp. 595–603.

Birnbaum, J. H., Wanner, D., Gietl, A. F., Saake, A., Kündig, T. M., Hock, C., Nitsch, R. M. and Tackenberg, C. (2018) ‘Oxidative stress and altered mitochondrial protein expression in the absence of amyloid-β and tau pathology in iPSC-derived neurons from sporadic Alzheimer’s disease patients’, Stem Cell Res, 27, pp. 121–130.

Button, R. W., Roberts, S. L., Willis, T. L., Hanemann, C. O. and Luo, S. (2017) ‘Accumulation of autophagosomes confers cytotoxicity’, J Biol Chem, 292(33), pp. 13599– 13614.

Capano, L. S., Sato, C., Ficulle, E., Yu, A., Horie, K., Kwon, J. S., Burbach, K. F., Barthélemy, N. R., Fox, S. G., Karch, C. M., Bateman, R. J., Houlden, H., Morimoto, R. I., Holtzman, D. M., Duff, K. E. and Yoo, A. S. (2022) ‘Recapitulation of endogenous 4R tau expression and formation of insoluble tau in directly reprogrammed human neurons’, Cell Stem Cell, 29(6), pp. 918–932.e8.

Cason, S. E., Mogre, S. S., Holzbaur, E. L. F. and Koslover, E. F. (2022) ‘Spatiotemporal analysis of axonal autophagosome-lysosome dynamics reveals limited fusion events and slow maturation’, Mol Biol Cell, 33(13), pp. ar123.

Cates, K., McCoy, M. J., Kwon, J. S., Liu, Y., Abernathy, D. G., Zhang, B., Liu, S., Gontarz, P., Kim, W. K., Chen, S., Kong, W., Ho, J. N., Burbach, K. F., Gabel, H. W., Morris, S. A. and Yoo, A. S. (2021) ‘Deconstructing Stepwise Fate Conversion of Human Fibroblasts to Neurons by MicroRNAs’, Cell Stem Cell, 28(1), pp. 127–140.e9.

Cates, K., Yuan, L., Yang, Y. and Yoo, A. S. (2025) ‘Fate erasure logic of gene networks underlying direct neuronal conversion of somatic cells by microRNAs’, Cell Rep, 44(1), pp. 115153.

Chakravorty, A., Jetto, C. T. and Manjithaya, R. (2019) ‘Dysfunctional Mitochondria and Mitophagy as Drivers of Alzheimer’s Disease Pathogenesis’, Front Aging Neurosci, 11, pp. 311.

Chen, E. Y., Tan, C. M., Kou, Y., Duan, Q., Wang, Z., Meirelles, G. V., Clark, N. R. and Ma’ayan, A. (2013) ‘Enrichr: interactive and collaborative HTML5 gene list enrichment analysis tool’, BMC Bioinformatics, 14, pp. 128.

Chou, C. C., Vest, R., Prado, M. A., Wilson-Grady, J., Paulo, J. A., Shibuya, Y., Moran-Losada, P., Lee, T. T., Luo, J., Gygi, S. P., Kelly, J. W., Finley, D., Wernig, M., Wyss-Coray, T. and Frydman, J. (2025) ‘Proteostasis and lysosomal repair deficits in transdifferentiated neurons of Alzheimer’s disease’, Nat Cell Biol, 27(4), pp. 619– 632.

Church, V. A., Cates, K., Capano, L., Aryal, S., Kim, W. K. and Yoo, A. S. (2021) ‘Generation of Human Neurons by microRNA-Mediated Direct Conversion of Dermal Fibroblasts’, Methods Mol Biol, 2239, pp. 77–100.

Cui, M., Tang, X., Christian, W. V., Yoon, Y. and Tieu, K. (2010) ‘Perturbations in mitochondrial dynamics induced by human mutant PINK1 can be rescued by the mitochondrial division inhibitor mdivi-1’, J Biol Chem, 285(15), pp. 11740–52.

Fang, E. F., Hou, Y., Palikaras, K., Adriaanse, B. A., Kerr, J. S., Yang, B., Lautrup, S., Hasan-Olive, M. M., Caponio, D., Dan, X., Rocktäschel, P., Croteau, D. L., Akbari, M., Greig, N. H., Fladby, T., Nilsen, H., Cader, M. Z., Mattson, M. P., Tavernarakis, N. and Bohr, V. A. (2019) ‘Mitophagy inhibits amyloid-β and tau pathology and reverses cognitive deficits in models of Alzheimer’s disease’, Nat Neurosci, 22(3), pp. 401– 412.

Farfel-Becker, T., Roney, J. C., Cheng, X. T., Li, S., Cuddy, S. R. and Sheng, Z. H. (2019) ‘Neuronal Soma-Derived Degradative Lysosomes Are Continuously Delivered to Distal Axons to Maintain Local Degradation Capacity’, Cell Rep, 28(1), pp. 51–64.e4.

Feng, Y. and Lu, Y. (2025) ‘The nuclear-mitochondrial crosstalk in aging: From mechanisms to therapeutics’, Free Radic Biol Med, 232, pp. 391–397.

Ferguson, S. M. (2019) ‘Neuronal lysosomes’, Neurosci Lett, 697, pp. 1–9.

Gan, X., Huang, S., Wu, L., Wang, Y., Hu, G., Li, G., Zhang, H., Yu, H., Swerdlow, R. H., Chen, J. X. and Yan, S. S. (2014) ‘Inhibition of ERK-DLP1 signaling and mitochondrial division alleviates mitochondrial dysfunction in Alzheimer’s disease cybrid cell’, Biochim Biophys Acta, 1842(2), pp. 220–31.

Grimm, A. and Eckert, A. (2017) ‘Brain aging and neurodegeneration: from a mitochondrial point of view’, J Neurochem, 143(4), pp. 418–431.

Harrison, D. E., Strong, R., Sharp, Z. D., Nelson, J. ., Astle, C. M., Flurkey, K., Nadon, N. L., Wilkinson, J. E., Frenkel, K., Carter, C. S., Pahor, M., Javors, M. A., Fernandez, E. and Miller, R. A. (2009) ‘Rapamycin fed late in life extends lifespan in genetically heterogeneous mice’, Nature, 460(7253), pp. 392–5.

Hawkins, K. E. and Duchen, M. (2019) ‘Modelling mitochondrial dysfunction in Alzheimer’s disease using human induced pluripotent stem cells’, World J Stem Cells, 11(5), pp. 236–253.

Herring, C. A., Simmons, R. K., Freytag, S., Poppe, D., Moffet, J. J. D., Pflueger, J., Buckberry, S., Vargas-Landin, D. B., Clément, O., Echeverría, E. ., Sutton, G. J., Alvarez-Franco, A., Hou, R., Pflueger, C., McDonald, K., Polo, J. M., Forrest, A. R. R., Nowak, A. K., Voineagu, I., Martelotto, L. and Lister, R. (2022) ‘Human prefrontal cortex gene regulatory dynamics from gestation to adulthood at single-cell resolution’, Cell, 185(23), pp. 4428–4447.e28.

Hidvegi, T., Ewing, M., Hale, P., Dippold, C., Beckett, C., Kemp, C., Maurice, N., Mukherjee, A., Goldbach, C., Watkins, S., Michalopoulos, G. and Perlmutter, D. H. (2010) ‘An autophagy-enhancing drug promotes degradation of mutant alpha1-antitrypsin Z and reduces hepatic fibrosis’, Science, 329(5988), pp. 229–32.

Horvath, S. (2013) ‘DNA methylation age of human tissues and cell types’, Genome Biol, 14(10), pp. R115.

Huh, C. J., Zhang, B., Victor, M. B., Dahiya, S., Batista, L. F., Horvath, S. and Yoo, A. S. (2016) ‘Maintenance of age in human neurons generated by microRNA-based neuronal conversion of fibroblasts’, Elife, 5.

Jeffries, A. M., Yu, T., Ziegenfuss, J. S., Tolles, A. K., Baer, C. E., Sotelo, C. B., Kim, Y., Weng, Z. and Lodato, M. A. (2025) ‘Single-cell transcriptomic and genomic changes in the ageing human brain’, Nature.

Juricic, P., Lu, Y. X., Leech, T., Drews, L. F., Paulitz, J., Lu, J., Nespital, T., Azami, S., Regan, J. C., Funk, E., Fröhlich, J., Grönke, S. and Partridge, L. (2022) ‘Long-lasting geroprotection from brief rapamycin treatment in early adulthood by persistently increased intestinal autophagy’, Nat Aging, 2(9), pp. 824–836.

Karch, C. M., Hernández, D., Wang, J. C., Marsh, J., Hewitt, A. W., Hsu, S., Norton, J., Levitch, D., Donahue, T., Sigurdson, W., Ghetti, B., Farlow, M., Chhatwal, J., Berman, S., Cruchaga, C., Morris, J. C., Bateman, R. J., Pébay, A., Goate, A. M. and (DIAN), D. I. A. N. (2018) ‘Human fibroblast and stem cell resource from the Dominantly Inherited Alzheimer Network’, Alzheimers Res Ther, 10(1), pp. 69.

Kaushik, S., Arias, E., Kwon, H., Lopez, N. M., Athonvarangkul, D., Sahu, S., Schwartz, G. J., Pessin, J. E. and Singh, R. (2012) ‘Loss of autophagy in hypothalamic POMC neurons impairs lipolysis’, EMBO Rep, 13(3), pp. 258–65.

Kim, K. B., Lee, S. and Kim, J. H. (2020) ‘Neuroprotective effects of urolithin A on H’, Nutr Res Pract, 14(1), pp. 3–11.

Kim, S. Y., Strucinska, K., Osei, B., Han, K., Kwon, S. K. and Lewis, T. L. (2022) ‘Neuronal mitochondrial morphology is significantly affected by both fixative and oxygen level during perfusion’, Front Mol Neurosci, 15, pp. 1042616.

Kim, Y., Zheng, X., Ansari, Z., Bunnell, M. C., Herdy, J. R., Traxler, L., Lee, H., Paquola, A. C. M., Blithikioti, C., Ku, M., Schlachetzki, J. C. M., Winkler, J., Edenhofer, F., Glass, C. K., Paucar, A. A., Jaeger, B. N., Pham, S., Boyer, L., Campbell, B. C., Hunter, T., Mertens, J. and Gage, F. H. (2018) ‘Mitochondrial Aging Defects Emerge in Directly Reprogrammed Human Neurons due to Their Metabolic Profile’, Cell Rep, 23(9), pp. 2550–2558.

Klee, C. B., Crouch, T. H. and Krinks, M. H. (1979) ‘Calcineurin: a calcium- and calmodulin-binding protein of the nervous system’, Proc Natl Acad Sci U S A, 76(12), pp. 6270–3.

Klinman, E. and Holzbaur, E. L. (2016) ‘Comparative analysis of axonal transport markers in primary mammalian neurons’, Methods Cell Biol, 131, pp. 409–24.

Kuleshov, M. V., Jones, M. R., Rouillard, A. D., Fernandez, N. F., Duan, Q., Wang, Z., Koplev, S., Jenkins, S. L., Jagodnik, K. M., Lachmann, A., McDermott, M. G., Monteiro, C. D., Gundersen, G. W. and Ma’ayan, A. (2016) ‘Enrichr: a comprehensive gene set enrichment analysis web server 2016 update’, Nucleic Acids Res, 44(W1), pp. W90–7.

Lapierre, L. R., Kumsta, C., Sandri, M., Ballabio, A. and Hansen, M. (2015) ‘Transcriptional and epigenetic regulation of autophagy in aging’, Autophagy, 11(6), pp. 867–80.

Lee, S. W., Oh, Y. M., Victor, M. B., Yang, Y., Chen, S., Strunilin, I., Dahiya, S., Dolle, R. E., Pak, S. C., Silverman, G. A., Perlmutter, D. H. and Yoo, A. S. (2024) ‘Longitudinal modeling of human neuronal aging reveals the contribution of the RCAN1-TFEB pathway to Huntington’s disease neurodegeneration’, Nat Aging, 4(1), pp. 95–109.

Lewis, T. L., Turi, G. F., Kwon, S. K., Losonczy, A. and Polleux, F. (2016) ‘Progressive Decrease of Mitochondrial Motility during Maturation of Cortical Axons In Vitro and In Vivo’, Curr Biol, 26(19), pp. 2602–2608.

Li, Y., Sheftic, S. R., Grigoriu, S., Schwieters, C. D., Page, R. and Peti, W. (2020) ‘The structure of the RCAN1:CN complex explains the inhibition of and substrate recruitment by calcineurin’, Sci Adv, 6(27).

Liang, C. S., Li, D. J., Yang, F. C., Tseng, P. T., Carvalho, A. F., Stubbs, B., Thompson, T., Mueller, C., Shin, J. I., Radua, J., Stewart, R., Rajji, T. K., Tu, Y. K., Chen, T. Y., Yeh, T. C., Tsai, C. K., Yu, C. L., Pan, C. C. and Chu, C. S. (2021) ‘Mortality rates in Alzheimer’s disease and non-Alzheimer’s dementias: a systematic review and meta-analysis’, Lancet Healthy Longev, 2(8), pp. e479–e488.

Lipinski, M. M., Zheng, B., Lu, T., Yan, Z., Py, B. F., Ng, A., Xavier, R. J., Li, C., Yankner, B. A., Scherzer, C. R. and Yuan, J. (2010) ‘Genome-wide analysis reveals mechanisms modulating autophagy in normal brain aging and in Alzheimer’s disease’, Proc Natl Acad Sci U S A, 107(32), pp. 14164–9.

Liu, Y. J., McIntyre, R. L., Janssens, G. E. and Houtkooper, R. H. (2020) ‘Mitochondrial fission and fusion: A dynamic role in aging and potential target for age-related disease’, Mech Ageing Dev, 186, pp. 111212.

Lu, Y. L., Liu, Y., McCoy, M. J. and Yoo, A. S. (2021) ‘MiR-124 synergism with ELAVL3 enhances target gene expression to promote neuronal maturity’, Proc Natl Acad Sci U S A, 118(22).

Luzio, J. P., Pryor, P. R. and Bright, N. A. (2007) ‘Lysosomes: fusion and function’, Nat Rev Mol Cell Biol, 8(8), pp. 622–32.

Maday, S., Wallace, K. E. and Holzbaur, E. L. (2012) ‘Autophagosomes initiate distally and mature during transport toward the cell soma in primary neurons’, J Cell Biol, 196(4), pp. 407–17.

Malik, B. R., Maddison, D. C., Smith, G. A. and Peters, O. M. (2019) ‘Autophagic and endo- lysosomal dysfunction in neurodegenerative disease’, Mol Brain, 12(1), pp. 100.

Martina, J. A., Chen, Y., Gucek, M. and Puertollano, R. (2012) ‘MTORC1 functions as a transcriptional regulator of autophagy by preventing nuclear transport of TFEB’, Autophagy, 8(6), pp. 903–14.

Mauvezin, C., Nagy, P., Juhász, G. and Neufeld, T. P. (2015) ‘Autophagosome-lysosome fusion is independent of V-ATPase-mediated acidification’, Nat Commun, 6, pp. 7007.

Mertens, J., Marchetto, M. C., Bardy, C. and Gage, F. H. (2016) ‘Evaluating cell reprogramming, differentiation and conversion technologies in neuroscience’, Nat Rev Neurosci, 17(7), pp. 424– 37.

Mertens, J., Paquola, A. C. M., Ku, M., Hatch, E., Böhnke, L., Ladjevardi, S., McGrath, S., Campbell, B., Lee, H., Herdy, J. R., Gonçalves, J. T., Toda, T., Kim, Y., Winkler, J., Yao, J., Hetzer, M. W. and Gage, F. H. (2015) ‘Directly Reprogrammed Human Neurons Retain Aging-Associated Transcriptomic Signatures and Reveal Age-Related Nucleocytoplasmic Defects’, Cell Stem Cell, 17(6), pp. 705–718.

Mertens, J., Reid, D., Lau, S., Kim, Y. and Gage, F. H. (2018) ‘Aging in a Dish: iPSC-Derived and Directly Induced Neurons for Studying Brain Aging and Age-Related Neurodegenerative Diseases’, Annu Rev Genet, 52, pp. 271–293.

Misgeld, T. and Schwarz, T. L. (2017) ‘Mitostasis in Neurons: Maintaining Mitochondria in an Extended Cellular Architecture’, Neuron, 96(3), pp. 651–666.

Mondal, S., Dubey, J., Awasthi, A., Sure, G. R., Vasudevan, A. and Koushika, S. P. (2021) ‘Tracking Mitochondrial Density and Positioning along a Growing Neuronal Process in Individual’, eNeuro, 8(4).

Nixon, R. A. (2020) ‘The aging lysosome: An essential catalyst for late-onset neurodegenerative diseases’, Biochim Biophys Acta Proteins Proteom, 1868(9), pp. 140443.

Nixon, R. A., Wegiel, J., Kumar, A., Yu, W. H., Peterhoff, C., Cataldo, A. and Cuervo, A. M. (2005) ‘Extensive involvement of autophagy in Alzheimer disease: an immuno-electron microscopy study’, J Neuropathol Exp Neurol, 64(2), pp. 113–22.

Oh, Y. M., Lee, S. W., Kim, W. K., Chen, S., Church, V. A., Cates, K., Li, T., Zhang, B., Dolle, R. ., Dahiya, S., Pak, S. C., Silverman, G. A., Perlmutter, D. H. and Yoo, A. S. (2022) ‘Age-related Huntington’s disease progression modeled in directly reprogrammed patient-derived striatal neurons highlights impaired autophagy’, Nat Neurosci, 25(11), pp. 1420–1433.

Oh, Y. M., Lee, S. W. and Yoo, A. S. (2023) ‘Modeling Huntington disease through microRNA-mediated neuronal reprogramming identifies age-associated autophagy dysfunction driving the onset of neurodegeneration’, Autophagy, 19(9), pp. 2613–2615.

Ott, C., König, J., Höhn, A., Jung, T. and Grune, T. (2016) Macroautophagy is impaired in old murine brain tissue as well as in senescent human fibroblasts’, Redox Biol, 10, pp. 266–273.

Palmieri, M., Impey, S., Kang, H., di Ronza, A., Pelz, C., Sardiello, M. and Ballabio, A. (2011) ‘Characterization of the CLEAR network reveals an integrated control of cellular clearance pathways’, Hum Mol Genet, 20(19), pp. 3852–66.

Pankiv, S., Clausen, T. H., Lamark, T., Brech, A., Bruun, J. A., Outzen, H., Øvervatn, A., Bjørkøy, G. and Johansen, T. (2007) ‘p62/SQSTM1 binds directly to Atg8/LC3 to facilitate degradation of ubiquitinated protein aggregates by autophagy’, J Biol Chem, 282(33), pp. 24131–45.

Patterson, M., Chan, D. N., Ha, I., Case, D., Cui, Y., Van Handel, B., Mikkola, H. K. and Lowry, W. E. (2012) ‘Defining the nature of human pluripotent stem cell progeny’, Cell Res, 22(1), pp. 178–93.

Pekkurnaz, G. and Wang, X. (2022) ‘Mitochondrial heterogeneity and homeostasis through the lens of a neuron’, Nat Metab, 4(7), pp. 802–812.

Pitrez, P. R., Monteiro, L. M., Borgogno, O., Nissan, X., Mertens, J. and Ferreira, L. (2024) ‘Cellular reprogramming as a tool to model human aging in a dish’, Nat Commun, 15(1), pp. 1816.

Rangaraju, V., Lauterbach, M. and Schuman, E. M. (2019) ‘Spatially Stable Mitochondrial Compartments Fuel Local Translation during Plasticity’, Cell, 176(1-2), pp. 73–84.e15.

Rangaraju, V., Lewis, T. L., Hirabayashi, Y., Bergami, M., Motori, E., Cartoni, R., Kwon, S. K. and Courchet, J. (2019) ‘Pleiotropic Mitochondria: The Influence of Mitochondria on Neuronal Development and Disease’, J Neurosci, 39(42), pp. 8200–8208.

Rusnak, F. and Mertz, P. (2000) ‘Calcineurin: form and function’, Physiol Rev, 80(4), pp. 1483–521.

Ryu, D., Mouchiroud, L., Andreux, P. A., Katsyuba, E., Moullan, N., Nicolet-Dit-Félix, A. A., Williams, E. G., Jha, P., Lo Sasso, G., Huzard, D., Aebischer, P., Sandi, C., Rinsch, C. and Auwerx, J. (2016) ‘Urolithin A induces mitophagy and prolongs lifespan in C. elegans and increases muscle function in rodents’, Nat Med, 22(8), pp. 879–88.

Sardiello, M., Palmieri, M., di Ronza, A., Medina, D. L., Valenza, M., Gennarino, V. A., Di Malta, C., Donaudy, F., Embrione, V., Polishchuk, R. S., Banfi, S., Parenti, G., Cattaneo, E. and Ballabio, A. (2009) ‘A gene network regulating lysosomal biogenesis and function’, Science, 325(5939), pp. 473–7.

Settembre, C., De Cegli, R., Mansueto, G., Saha, P. K., Vetrini, F., Visvikis, O., Huynh, T., Carissimo, A., Palmer, D., Klisch, T. J., Wollenberg, A. C., Di Bernardo, D., Chan, L., Irazoqui, J. E. and Ballabio, A. (2013) ‘TFEB controls cellular lipid metabolism through a starvation-induced autoregulatory loop’, Nat Cell Biol, 15(6), pp. 647–58.

Settembre, C., Di Malta, C., Polito, V. A., Garcia Arencibia, M., Vetrini, F., Erdin, S., Erdin, S. U., Huynh, T., Medina, D., Colella, P., Sardiello, M., Rubinsztein, D. C. and Ballabio, A. (2011) ‘TFEB links autophagy to lysosomal biogenesis’, Science, 332(6036), pp. 1429–33.

Stavoe, A. K. H. and Holzbaur, E. L. F. (2020) ‘Neuronal autophagy declines substantially with age and is rescued by overexpression of WIPI2’, Autophagy, 16(2), pp. 371–372.

Sun, Z., Kwon, J. S., Ren, Y., Chen, S., Walker, C. K., Lu, X., Cates, K., Karahan, H., Sviben, S., Fitzpatrick, J. A. J., Valdez, C., Houlden, H., Karch, C. M., Bateman, R. J., Sato, C., Mennerick, S. ., Diamond, M. I., Kim, J., Tanzi, R. E., Holtzman, D. M. and Yoo, A. S. (2024) ‘Modeling late-onset Alzheimer’s disease neuropathology via direct neuronal reprogramming’, Science, 385(6708), pp. adl2992.

Sure, G. R., Chatterjee, A., Mishra, N., Sabharwal, V., Devireddy, S., Awasthi, A., Mohan, S. and Koushika, S. P. (2018) ‘UNC-16/JIP3 and UNC- 76/FEZ1 limit the density of mitochondria in C. elegans neurons by maintaining the balance of anterograde and retrograde mitochondrial transport’, Sci Rep, 8(1), pp. 8938.

Swerdlow, R. H. (2018) ‘Mitochondria and Mitochondrial Cascades in Alzheimer’s Disease’, J Alzheimers Dis, 62(3), pp. 1403–1416.

Tian, Y. E., Cropley, V., Maier, A. B., Lautenschlager, N. T., Breakspear, M. and Zalesky, A. (2023) ‘Heterogeneous aging across multiple organ systems and prediction of chronic disease and mortality’, Nat Med, 29(5), pp. 1221– 1231.

Toescu, E. C., Myronova, N. and Verkhratsky, A. (2000) ‘Age-related structural and functional changes of brain mitochondria’, Cell Calcium, 28(5-6), pp. 329–38.

Tsong, H., Holzbaur, E. L. and Stavoe, A. K. (2023) ‘Aging Differentially Affects Axonal Autophagosome Formation and Maturation’, Autophagy, 19(12), pp. 3079–3095.

Varghese, N., Szabo, L., Cader, M. Z., Lejri, I., Grimm, A. and Eckert, A. (2025) ‘Tracing mitochondrial marks of neuronal aging in iPSCs-derived neurons and directly converted neurons’, Commun Biol, 8(1), pp. 723.

Victor, M. B., Richner, M., Hermanstyne, T. O., Ransdell, J. L., Sobieski, C., Deng, P. Y., Klyachko, V. A., Nerbonne, J. M. and Yoo, A. S. (2014) ‘Generation of human striatal neurons by microRNA-dependent direct conversion of fibroblasts’, Neuron, 84(2), pp. 311–23.

Victor, M. B., Richner, M., Olsen, H. E., Lee, S. W., Monteys, A. M., Ma, C., Huh, C. J., Zhang, B., Davidson, B. L., Yang, X. W. and Yoo, A. S. (2018) ‘Striatal neurons directly converted from Huntington’s disease patient fibroblasts recapitulate age-associated disease phenotypes’, Nat Neurosci, 21(3), pp. 341–352.

Wang, Y., Cobanoglu, M. C., Li, J., Hidvegi, T., Hale, P., Ewing, M., Chu, A. S., Gong, Z., Muzumdar, R., Pak, S. C., Silverman, G. A., Bahar, I. and Perlmutter, D. H. (2019) ‘An analog of glibenclamide selectively enhances autophagic degradation of misfolded α1-antitrypsin Z’, PLoS One, 14(1), pp. e0209748.

Wareski, P., Vaarmann, A., Choubey, V., Safiulina, D., Liiv, J., Kuum, M. and Kaasik, A. (2009) ‘PGC-1{alpha} and PGC-1{beta} regulate mitochondrial density in neurons’, J Biol Chem, 284(32), pp. 21379–85.

Wu, Y. Y., Chiu, F. L., Yeh, C. S. and Kuo, H. C. (2019) ‘Opportunities and challenges for the use of induced pluripotent stem cells in modelling neurodegenerative disease’, Open Biol, 9(1), pp. 180177.

Xie, Z., Bailey, A., Kuleshov, M. V., Clarke, D. J. B., Evangelista, J. E., Jenkins, S. L., Lachmann, A., Wojciechowicz, M. L., Kropiwnicki, E., Jagodnik, K. M., Jeon, M. and Ma’ayan, A. (2021) ‘Gene Set Knowledge Discovery with Enrichr’, Curr Protoc, 1(3), pp. e90.

Xu, F., Armstrong, R., Urrego, D., Qazzaz, M., Pehar, M., Armstrong, J. N., Shutt, T. and Syed, N. (2016) ‘The mitochondrial division inhibitor Mdivi-1 rescues mammalian neurons from anesthetic-induced cytotoxicity’, Mol Brain, 9, pp. 35.

Yoo, A. S., Sun, A. X., Li, L., Shcheglovitov, A., Portmann, T., Li, Y., Lee-Messer, C., Dolmetsch, R. E., Tsien, R. W. and Crabtree, G. R. (2011) ‘MicroRNA-mediated conversion of human fibroblasts to neurons’, Nature, 476(7359), pp. 228–31.

Zacharioudakis, E., Agianian, B., Kumar Mv, V., Biris, N., Garner, T. P., Rabinovich-Nikitin, I., Ouchida, A. T., Margulets, V., Nordstrøm, L. U., Riley, J. S., Dolgalev, I., Chen, Y., Wittig, A. J. H., Pekson, R., Mathew, C., Wei, P., Tsirigos, A., Tait, S. W. G., Kirshenbaum, L. A., Kitsis, R. N. and Gavathiotis, E. (2022) ‘Modulating mitofusins to control mitochondrial function and signaling’, Nat Commun, 13(1), pp. 3775.

Zhang, B., Zhang, X., Treebupachatsakul, W., Pantan, R., Kampan, N., Phatsara, M., Shi, C. and Narakornsak, S. (2025) ‘Neuroprotective effect of Urolithin A via downregulating VDAC1-mediated autophagy in Alzheimer’s disease’, Acta Histochem, 127(4), pp. 152290.

Zhang, Y., Lanjuin, A., Chowdhury, S. R., Mistry, M., Silva-García, C. G., Weir, H. J., Lee, C. L., Escoubas, C. C., Tabakovic, E. and Mair, W. B. (2019) ‘Neuronal TORC1 modulates longevity via AMPK and cell nonautonomous regulation of mitochondrial dynamics in’, Elife, 8.

